# Discovery and biosynthesis of persiathiacins: Unusual polyglycosylated thiopeptides active against multi-drug resistant tuberculosis

**DOI:** 10.1101/2021.10.24.465558

**Authors:** Yousef Dashti, Fatemeh Mohammadipanah, Matthew Belousoff, Anthony Vocat, Daniel Zabala, Christopher D Fage, Isolda Romero-Canelon, Boyke Bunk, Cathrin Spröer, Jörg Overmann, Stewart T Cole, Gregory L Challis

**Author notes:** Y.D. The Centre for Bacterial Cell Biology, Biosciences Institute, Medical School, Newcastle University, Newcastle upon Tyne, NE2 4AX, UK; S.T.C. Institut Pasteur, 25 – 28 rue du Docteur Roux, Paris, France.

## Abstract

Thiopeptides are ribosomally biosynthesized and post-translationally modified peptides (RiPPs) that potently inhibit the growth of Gram-positive bacteria by targeting multiple steps in protein biosynthesis. The poor pharmacological properties of thiopeptides, in particular their low aqueous solubility, has hindered their development into clinically useful antibiotics. Antimicrobial activity screens of a library of Actinobacterial extracts led to discovery of the novel polyglycosylated thiopeptides persiathiacins A and B from *Actinokineospora* sp. UTMC 2475 and *Actinokineospora* sp. UTMC 2448. Persiathiacin A is active against methicillin-resistant *Staphylococcus aureus* (MRSA) and several *Mycobacterium tuberculosis* strains, including drug-resistant and multidrug-resistant clinical isolates, and does not significantly affect the growth of ovarian cancer cells at concentrations up to 400 μM. *In vitro* translation assays showed that, like other thiopeptide antibiotics, persiathiacin A targets protein biosynthesis. Polyglycosylated thiopeptides are extremely rare and nothing is known about their biosynthesis. Sequencing and analysis of the *Actinokineospora* sp. UTMC 2448 genome enabled identification of the putative persiathiacin biosynthetic gene cluster. A cytochrome P450 encoded by this gene cluster catalyses the hydroxylation of nosiheptide *in vitro* and *in vivo*, consistent with the proposal that the cluster directs persiathiacin biosynthesis. Several genes in the cluster encode homologues of enzymes known to catalyse the assembly and attachment of deoxysugars during the biosynthesis of other classes of glycosylated natural products. The discovery of the persiathiacins and their biosynthetic gene cluster thus provides the basis for the development of biosynthetic engineering approaches to the creation of novel (poly)glycosylated thiopeptide derivatives with enhanced pharmacological properties.

## Introduction

During the past decade, *Mycobacterium tuberculosis* has caused about 30 million deaths worldwide, making it the second most deadly pathogen after the human immunodeficiency virus (HIV).^1^ Currently, 6-12 month multidrug regimens are prescribed to treat *M. tuberculosis* infections. However, due to difficulties with dosing, side effects, and the emergence of multi- and extensively drug-resistant strains, more effective antibiotics must be developed to combat this critical-priority pathogen.^2, 3^

Thiopeptide antibiotics are ribosomally biosynthesized and post-translationally modified peptides (RiPPs). They are assembled from ribosomal peptide precursors via an extensive array of post-translational modifications catalysed by a series of diverse enzymes.^4, 5^ The precursor peptides consist of an N-terminal leader region that acts as a recognition motif for most of the post-translational modification enzymes and a C-terminal core region that is incorporated into the mature product(s).^5, 6^ Common post-translational modifications of thiopeptides include azole formation via cyclodehydration/oxidation, dehydration of selected serine and threonine residues, and macrocyclization via a [4+2] cycloaddition. Some thiopeptides are further modified via the introduction of additional macrocycles or the attachment of hydroxyl, methyl, indolyl, or quinaldyl substituents to the core peptide.^7–13^ Many thiopeptides possess potent activity against clinically relevant bacteria, in addition to antitumor and immunosuppressive properties.^14^ For instance, nosiheptide **1**, nocathiacin I **2**, and philipimycin **3**, which are representative of “series e” thiopeptides, are active against MRSA and/or clinical isolates of *M. tuberculosis* (Figure 1).^15–17^ Despite their promising bioactivity, thiopeptides have failed to reach the clinic due to poor aqueous solubility and gastrointestinal absorption. Several strategies, including biosynthetic pathway engineering, analogue total synthesis, and semisynthetic modification, have been applied to produce analogues with improved pharmacological properties.^18^

**Figure 1.**
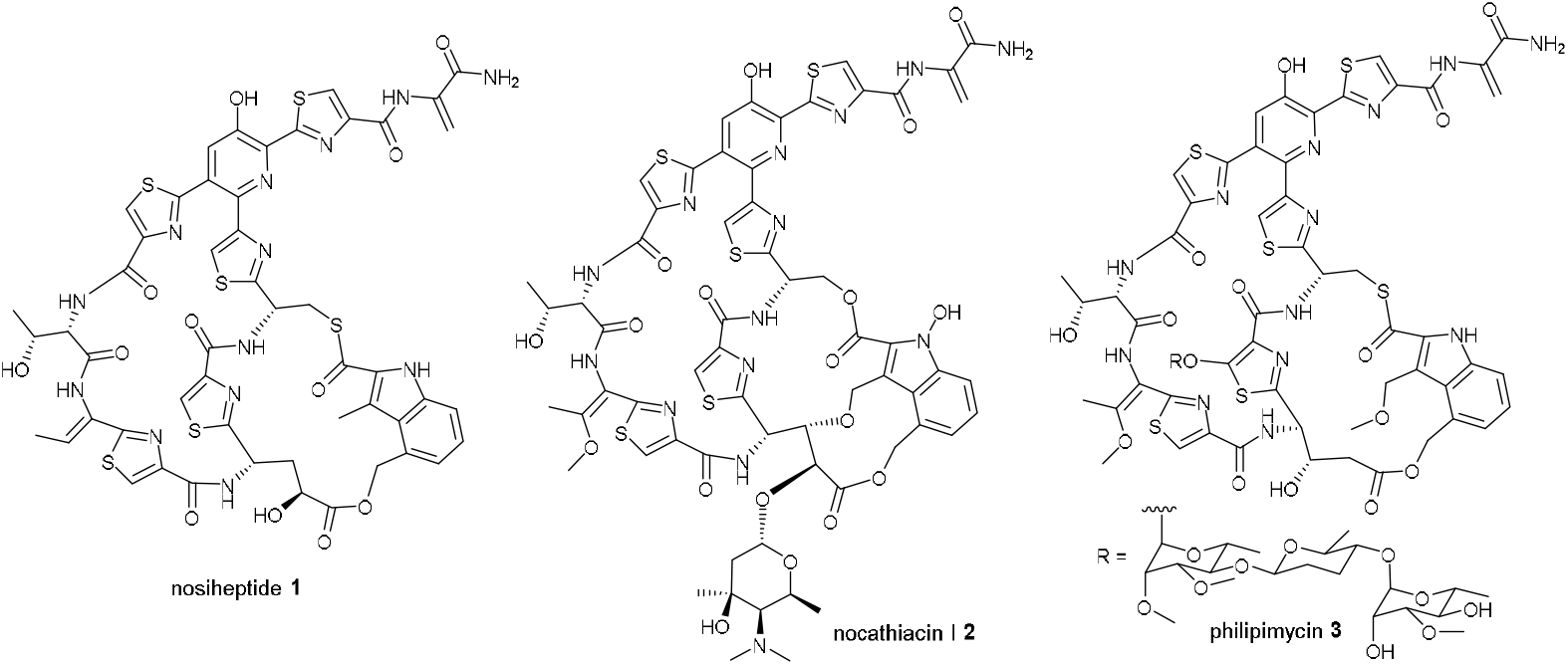
Structures of nosiheptide **1**, nocathiacin I **2** and philipimycin **3**, which are examples of “series e” thiopeptide antibiotics.

Here, we report the discovery of the novel polyglycosylated thiopeptide antibiotics persiathiacins A and B, which are active against methicillin-resistant *Staphylococcus aureus* (MRSA) and drug-resistant *M. tuberculosis* clinical isolates. The persiathiacins are the first example of naturally occurring thiopeptides with a glycosylated hydroxypyridine and only the second example of antibiotics belonging to this class bearing a polyglycosylated hydroxythiazole. Glycosylation of the hydroxypyridine in nocathiacin has been reported to significantly improve aqueous solubility and, more generally, glycosylation is a widely used strategy for increasing the solubility and circulatory half-life of therapeutic peptides.^19–21^ Thus, the discovery of the persiathiacins and the gene cluster directing their biosynthesis provides new opportunities for the development of biosynthetic engineering strategies for the creation of novel (poly)glycosylated thiopeptides with improved pharmacological properties.

## Results and discussion

### Isolation and structure elucidation of persiathiacins A and B

During a search for novel natural products with activity against MRSA, Actinobacteria isolated from various locations in Iran were screened for antibiotic production. Ethyl acetate extracts of *Actinokineospora* sp. UTMC 2475 and *Actinokineospora* sp. UTMC 2448 were found to exhibit potent activity against MRSA. The metabolite profiles of the extracts from these two strains were found to be very similar in UHPLC-ESI-Q-TOF-MS analyses. To identify the active metabolite(s), *Actinokineospora* sp. UTMC 2448 was cultured on solid ISP2 medium for 7 days, followed by ethyl acetate extraction and fractionation by semipreparative HPLC. A molecular formula of C_80_H_91_N_13_O_30_S_5_ was established from positive ion mode HR-ESI-MS and NMR data for the metabolite purified from the MRSA-active fraction. The planar structure of this compound, which we named persiathiacin A **4**, was elucidated using 1D and 2D NMR experiments (Figure 2, Figures S1 and S3-S8, Table S1). Characteristic signals for amino acid alpha protons at *δ_H_* 4.21, 5.62, and 5.80, which correlated in HSQC spectra with alpha carbon signals at *δ_C_* 56.2, 48.6, and 48.7 and in COSY spectra with signals for exchangeable amide protons at *δ_H_* 7.87, 7.89, and 8.24, indicated that the structure contains several amino acid residues. Four disubstituted thiazoles (including three bearing an acyl substituent at C-4) and a tetrasubstituted pyridine were identified based on distinctive singlets due to protons attached to sp^2^ hybridised carbons in the ^1^H NMR spectrum, the chemical shifts of the signals for the directly connected carbon atoms, and HMBC correlations between these protons and neighbouring carbons. A characteristic signal due to the sp^2^ hybridised methylene carbon of dehydroalanine (Dha) at *δ_C_* 104.8, which showed HSQC correlations to two protons at *δ_H_* 5.53 and 6.47, was also observed in the ^13^C NMR spectrum. This was further confirmed through ^*2J*^HMBC correlations between the methylene protons and the quaternary α-carbon of Dha at *δ_C_* 133.5 and a ^*3J*^ correlation to the carbonyl carbon at *δ_C_* 166.5. Taken together, these data indicated that persiathiacin A has a thiopeptide core structure.

**Figure 2.**
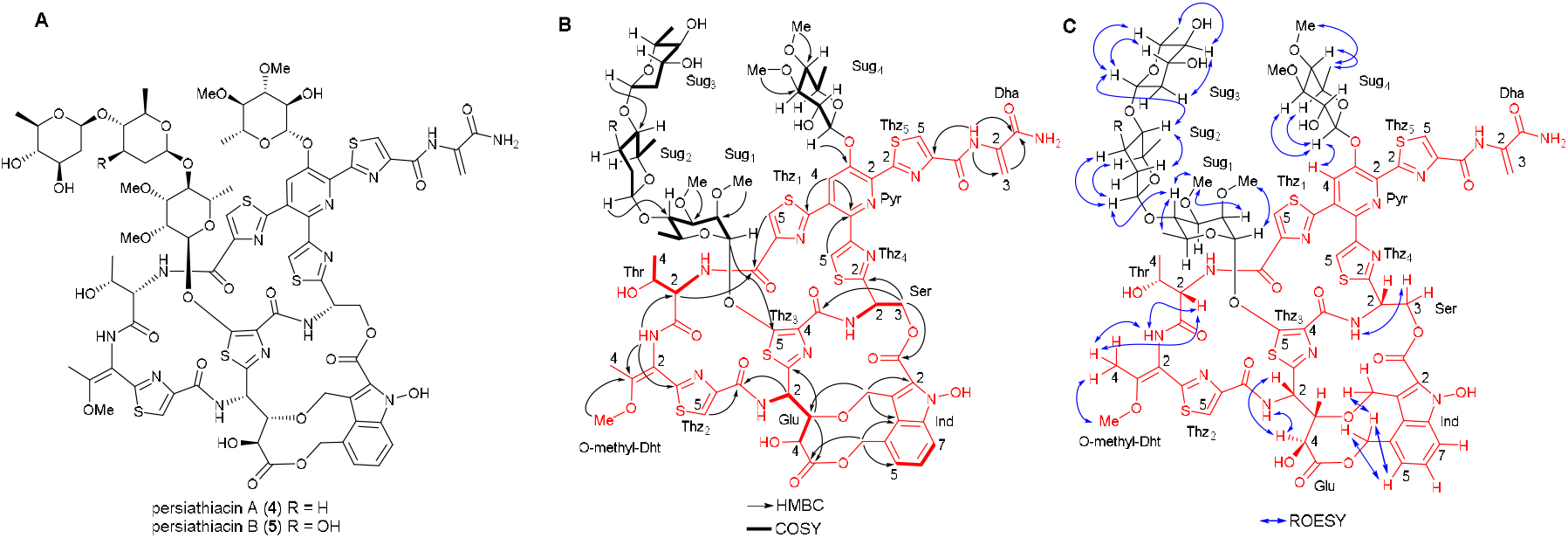
(A) Planar structures of persiathiacins A **4** and B **5**. (B) Summary of COSY and key HMBC correlations used to assign the planar structure of persiathiacins A **4** and B **5**. (C) Summary of key ROESY correlations observed for persiathiacins A **4** and B **5**.

The following HMBC data showed that the core thiopeptide is very similar to that of the nocathiacins^22^: A ^*3J*^ correlation between the O-methyl protons (*δ_H_* 3.78) and C3 (*δ_C_* 158.7) of the O-methyl-dehydrothreonine (O-methyl-Dht) residue; ^*3J*^ correlations between one of the methylene protons (*δ_H_* 4.17) in the 3-alkoxymethyl substituent of the indole and C3 (*δ_C_* 82.9) of the Glu residue, and between the C3 methine proton (*δ_H_* 3.65) of the Glu residue and the methylene carbon (*δ_C_* 65.9) of the indole 3-alkoxymethyl substituent; and a ^*3J*^ correlation between an exchangeable hydroxyl group proton (*δ_H_* 10.46) and C2 (*δ_C_* 127.0) of the indole. The linkage of the 2-carboxyl group of the indole to the side chain of the serine residue was identified through the distinctive chemical shift of the signal due to the carbonyl carbon (*δ_C_* 161.4) in comparison to those reported for nocathiacin I (161.1 ppm in DMSO-*d_6_*) and nosiheptide (181.80 ppm in DMSO-*d_6_*) as well as the chemical shift of the signal for C3 of the Ser residue (*δ_C_* 64.5) in comparison to the values reported for the corresponding carbon in nocathiacin I (63.3 ppm in DMSO-*d_6_*) and nosiheptide (29.5 ppm in DMSO-*d_6_*).^22, 23^ A ROESY correlation between the protons attached to C4 and the NH of the 4-O-methyl-Dht residue established that it contains an *E*-configured double bond.

Studies of the solution conformation of nocathiacin I, indicate that the amide proton of the 4-O-methyl-Dht residue and the α-proton of the Thr residue, the amide proton and the γ-proton of the Glu residue, and the α and γ-protons of the Glu residue, respectively, are in close spatial proximity.^24^ ROESY correlations between the corresponding protons in persiathiacin A are consistent with the Thr α-carbon and the stereocentres in the Glu residue having the same relative configurations as in nocathiacin I. Similarly, the splitting pattern of the signal for the less shielded of the diastereotopic C3 protons in the Ser residue, and a ROESY correlation between this proton and the Ser amide proton, indicate that the Ser α-carbon has the same relative configuration as in nocathiacin I. The only ambiguities are the relative stereochemistry of the β-carbon of the Thr residue and the absolute configuration of persiathiacin A. Given that persiathiacin A derives from a ribosomally-biosynthesised precursor and there is a high degree of similarity between the persiathicin and nocathiacin biosynthetic gene clusters (see below), it seems highly likely that the Thr residue has the L, rather than the L-*allo*, D, or D-*allo*, configuration, as reported for nocathiacin I.^24^

In addition to the resonances assigned to the core thiopeptide, four distinctive signals at *δ_C_* 100.4, 101.1, 101.9, and 102.2, assignable to anomeric carbons, that correlate in the HSQC spectrum with anomeric proton resonances at *δ_H_* 5.18, 4.43, 5.41, and 4.61, respectively, were observed. These indicated that the thiopeptide core is decorated with four glycosyl residues (sugars 1-4). Four independent coupled proton spin systems, indicative of four distinct 6-deoxysugars, were identified by analysis of the COSY and HMBC spectra, and *^3^J_HH_* coupling constants (Figure 2 and Table S1). The HMBC spectrum was also used to identify the attachment site of each sugar and the locations of four O-methyl groups. A correlation between the anomeric proton of sugar 1 (*δ_H_* 5.41) and C5 of thiazole 3 (*δ_C_* 160.3) showed that sugar 1 is attached to C5 of thiazole 3. The positions of the O-methyl groups in sugar 1 were assigned based on 3-bond correlations between the protons of the methoxy groups and the carbons they are attached to. Thus, the protons in one of the O-methyl groups (*δ_H_* 3.48) correlated with C2 (*δ_C_* 76.0), while the protons in the other O-methyl group (*δ_H_* 3.42) correlated with C3 (*δ_C_* 80.1). HMBC correlations between the anomeric proton of sugar 2 (*δ_H_* 4.61) and C4 of sugar 1 (*δ_C_* 77.0), and between the C4 proton (*δ_H_* 3.46) of sugar 1 and the anomeric carbon of sugar 2 (*δ_C_* 102.2) established the connectivity between sugars 1 and 2. Similarly, HMBC correlations between the anomeric proton of sugar 3 (*δ_H_* 4.43) and C4 of sugar 2 (*δ_C_* 80.60), and between the C4 proton of sugar 2 (*δ_H_* 3.08) and the anomeric carbon of sugar 3 (*δ_C_* 101.1) showed that sugar 3 is attached to the C4 hydroxyl group of sugar 2. An HMBC correlation between the anomeric proton of sugar 4 (*δ_H_* 5.18) and C3 of the pyridine (*δ_C_* 149.0), in addition to a ROESY correlation between the C1 proton of sugar 4 and the C4 proton of the pyridine (*δ_H_* 7.77), were consistent with the attachment of sugar 4 to the C3 hydroxyl group of the pyridine. Finally, HMBC correlations between the protons in one of the methoxy groups (*δ_H_* 3.45) and C3 (*δ_C_* 84.2) and the other methoxy group (*δ_H_* 3.51) and C4 (*δ_C_* 77.6) established the location of the O-methyl groups in sugar 4.

The signal due to the anomeric proton of sugar 1 is a broad singlet, suggesting it is the α-anomer, whereas the corresponding signals for sugars 2, 3 and 4 are doublets with ^3^*J_HH_* values of 9.0, 10.0 and 7.5 Hz, respectively, indicative of β-anomeric linkages. Moreover, these coupling constants indicate that the protons attached to C2 of sugars 2, 3, and 4 are all axial. ROESY correlations between the C1 proton and the C2 methoxy group, the C2 and C4 protons and the C3 methoxy group and the C4 and C6 protons in sugar 1 are consistent with this being a 2, 3-di-O-methyl-α-L-rhamnose, as reported for the corresponding sugar in the philipimycins.^17^ Similarly, ROESY correlations between the protons attached to C1 and C5, and C2 and C4 in sugar 2 suggests they are all axial, consistent with this being D-amecitose, as also observed in the philipimycins. Sugar 3 is assigned as D-olivose, based on ROESY correlations between the protons attached to C1 and C3, C1 and C5, and C2 and C4. Finally, ^3^*J*_HH_ values of 7.5 and 10.0 Hz for the proton attached to C2, and ROESY correlations between the protons attached to C1 and C3, and C1 and C5, indicate that H1, H2, H3 and H5 in sugar 4 are all axial. Given that the persiathiacin biosynthetic gene cluster encodes only a single NDP-hexose 4-ketoreductase, which is required for the biosynthesis of D-amecitose and D-olivose (see below), both of which have an axial C4 proton, we propose that sugar 4 is a 3, 4-di-O-methyl-6-deoxy-β-D-glucose residue.

In addition to persiathiacin A, a minor metabolite with a mass 16 Da greater than that of persiathiacin A was purified from the MRSA-active fractions of the *Actinokineospora* sp. UTMC 2448 extract. The molecular formula of this metabolite was deduced to be C_80_H_91_N_13_O_31_S_5_ from positive ion made HR-ESI-MS and NMR spectroscopic data, indicating that it is a persiathiacin A derivative containing an additional oxygen atom. The NMR spectroscopic data for this minor compound, which we named persiathiacin B **5**, was almost identical to that for persiathiacin A, except for the proton and carbon resonances of sugar 2 (Figures S2 and S9-S14, Table S2). Detailed analysis of 1D and 2D NMR spectra indicated that the β-D-amicetose residue in persiathiacin A is replaced by β-D-olivose in persiathiacin B (Figure 2).

### Identification and analysis of the putative persiathiacin biosynthetic gene cluster

To identify the persiathiacin biosynthetic gene cluster (BGC), the genome of *Actinokineospora* sp. UTMC 2448 was sequenced using single molecule real time (SMRT) sequencing. A complete circular genome sequence consisting of 7,012,397 bp was obtained using this approach (GenBank accession number CP031087). Analysis of the sequence using antiSMASH identified 32 putative specialized metabolite BGCs (Table S3).^25^ Among these, a cluster containing 33 genes (cluster 11; Table S3), several of which encode homologues of enzymes involved in the biosynthesis of other thiopeptides, was postulated to direct persiathiacin biosynthesis (Figure 3). Sequence comparisons showed that the products of *perA-perP* have a significant degree of similarity to the proteins encoded by *nosA-nosP* and *nocA-nocP* in the nosiheptide and nocathiacin BGCs (Table S4). Homologues of five additional genes in the nocathiacin BGC (*nocR* and *nocT-nocV*) that are absent from the nosiheptide cluster, are present in the putative persiathiacin cluster (*perR* and *perT-perV*, respectively; Figure 3).

**Figure 3.**
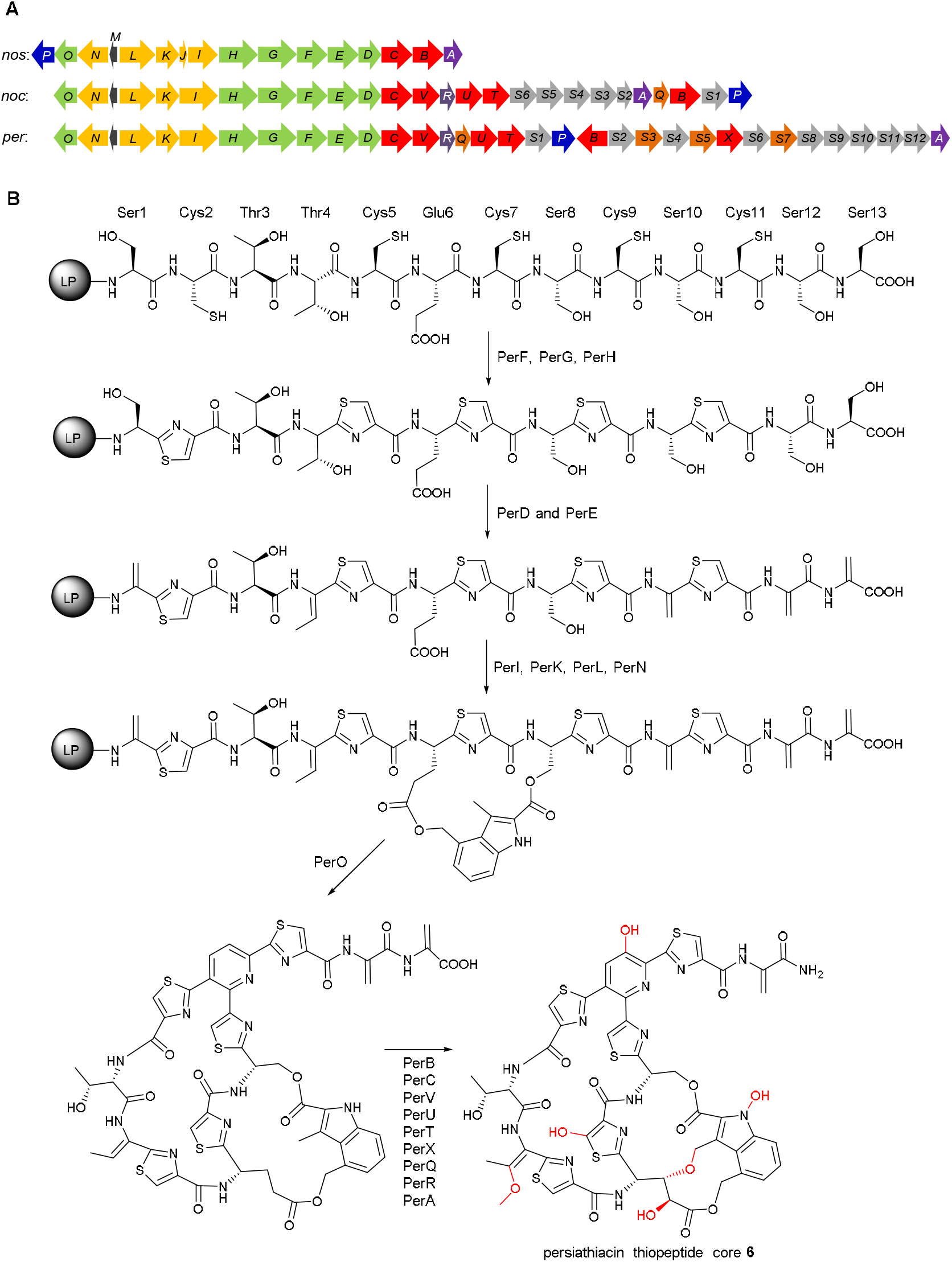
(A) Comparison of the nosiheptide, nocathiacin, and putative persiathiacin biosynthetic gene clusters. Genes are coloured as follows. Green: thiazole/pyridine formation and Ser/Thr dehydration; yellow: 3,4-dimethylindolic acid formation and attachment; red; cytochromes P450; grey: 6-deoxysugar biosynthesis and attachment; brown: methyltransferases. (B) The biosynthesis of the thiopeptide core **6** of the persiathiacins is proposed to commence with transcription and translation of *perM* to yield a precursor peptide comprised of an N-terminal leader peptide (LP) fused to a C-terminal core peptide (structure depicted). The core peptide undergoes a series of posttranslational modifications catalysed by several enzymes encoded by the persiathiacin biosynthetic gene cluster. See main text for further details.

Moreover, the putative persiathiacin BGC contains twelve genes (*perS1-perS12*) hypothesised to be responsible for the biosynthesis and attachment of four 6-deoxysugars to the thiopeptide core (Table S4).

Detailed sequence analysis of *perA-perV* and *perS1-perS12* enabled us to propose a biosynthetic pathway for persiathiacins A and B (Figures 3 and 5). First, *perM* is transcribed and translated into a 49 amino acid (aa) precursor peptide, consisting of a 36 aa N-terminal leader peptide fused to a 13 aa C-terminal core peptide with the sequence SCTTCECSCSCSS, which is fully consistent with the thiopeptide core structure of the persiathiacins deduced from the spectroscopic data.

**Figure 4.**
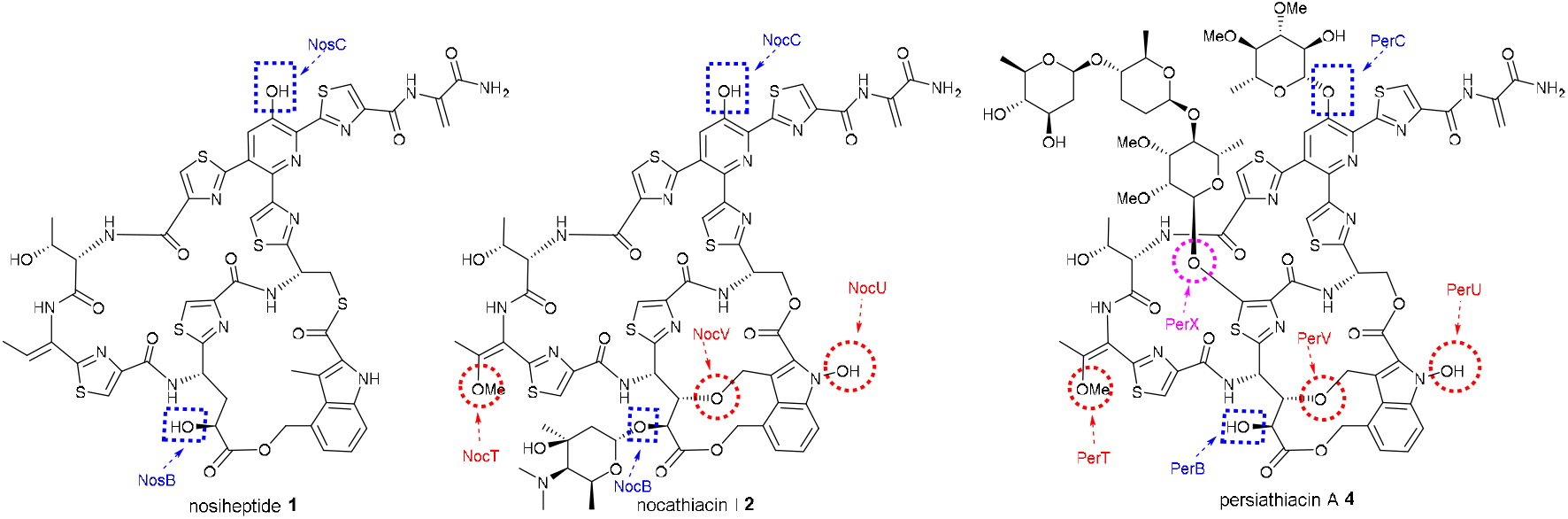
Comparative analysis of the structures of nosiheptide **1**, nocathiacin I **2**, and persiathiacin A **4** and the functions of the CYPs encoded by their BGCs. Blue dashed boxes highlight hydroxyl groups proposed to be installed by homologous CYPs (NosB/NocB/PerB and NosC/NocC/PerC) encoded by all three BGCs. Red dashed circles highlight hydroxyl groups proposed to be introduced by CYPs (NocT/PerT, NocV/PerV, and NocU/PerU) encoded by the nocathiacin and persiathiacin BGCs, but not the nosiheptide BGC. A purple dashed circle highlights the hydroxyl group proposed to be introduced by The CYP encoded by *perX*, which is only present in the persiathiacin BGC.

Cyclodehydration of the cysteine residues is proposed to be catalysed by PerG and PerH, followed by dehydrogenation catalysed by PerF to yield the five thiazoles in the persiathiacins.^7^ Putative dehydratases PerD and PerE are proposed to further modify the core peptide by catalysing selective dehydration of Ser1, Ser10, Ser12, Ser13, and Thr4.

Four enzymes encoded by *perI*, *perK*, *perL*, and *perN* are proposed to be responsible for the production of 3,4-dimethylindolic acid (DMIA) from L-tryptophan and its attachment to the core peptide (Figures 3 and S14). PerL is a putative radical *S*-adenosylmethionine (SAM) enzyme that is homologous to NosL, which transforms L-tryptophan into 3-methyl-2-indolic acid (MIA) via unusual constitutional isomerisation of a C-C bond.^26–34^ In nosiheptide biosynthesis, the ATP-dependent NosI enzyme adenylates MIA and loads it onto the phosphopantetheine thiol of the acyl carrier protein (ACP) NosJ. NosK then transfers the MIA residue to Cys8 of the nosiheptide core peptide (Figure S15).^35–37^ The persiathiacin and nocathiacin BGCs both lack *nosJ* homologues. Sequence comparisons of NosJ with PerI and PerK revealed similarity between NosJ and the C-terminus of the PerK. Similarly, it has previously been noted that the C-terminus of NocK is homologous to NosJ.^35^ Thus, it appears that in persiathiacin and nocathiacin biosynthesis, the C-terminal ACP domains of PerK and NocK are loaded with MIA by PerI and NocI, respectively. The N-terminal domains of PerK and NocK then catalyse attachment of the MIA residue to Ser8 of the persiathiacin and nocathiacin core peptides, respectively (Figure S14). Subsequently, the putative radical SAM methylase, PerN, is proposed, by analogy with the well-characterised mechanism of NosN,^35^ to catalyse methylenation of C4 in the MIA residue. The resulting electrophilic intermediate is attacked by the Glu6 carboxylate to form an ester linkage.^38^ Finally, PerO, which has >50% sequence identity to NosO and NocO, is hypothesised to be responsible for formation of the macrocycle and pyridine in the persiathiacins via a [4+2] cycloaddition.^39, 40^

Of the six putative cytochromes P450 (CYPs) encoded by the persiathiacin BGC, two (PerB and PerC) are homologous to NosB/NocB and NosC/NocC, which hydroxylate C3 of Glu6 and the pyridine, respectively.^13^ The CYPs encoded by *perV*, *perU*, and *perT* are similar in sequence to NocV, NocU, and NocT, respectively, encoded by the nocathiacin BGC. Genes encoding homologues of these enzymes are absent from the nosiheptide BGC. PerV is proposed to perform an analogous function to NocV - formation of the ether linkage between the indole and core peptide via a mechanism yet to be identified.^41^ Similarly, PerU is hypothesised to catalyse *N*-hydroxylation of the indole, by analogy with the proposed function of NocU.^42^ Comparison of the structures of persiathiacin A, nocathiacin I and nosiheptide (Figure 4) suggests the putative CYPs encoded by *perT*/*nocT* and methyltransferases encoded *perQ*/*nocQ* catalyse hydroxylation and subsequent O-methylation of the dehydrobutyrine residue to form the corresponding O-methyl-Dht residue. The only CYP-encoding gene in the persiathiacin BGC that does not have a homologue in either the nosiheptide or nocathiacin BGCs is *perX*. Structural comparison of persiathiacin A with nocathiacin I and nosiheptide suggests that the enzyme encoded by this gene catalyses hydroxylation of C5 in thiazole 3, to create the attachment site for the trisaccharide (Figure 4).

The final enzyme proposed to be involved in the assembly of the thiopeptide core of the persiathiacins is PerA. This is homologous to NosA, which catalyses dealkylative cleavage of the C-terminal Dha residue in nosiheptide biosynthesis, resulting in formation of the corresponding amide.^43^ PerA is proposed to catalyse an analogous reaction in persiathiacin biosynthesis (Figure 4).

The enzymes encoded by *perS1-perS12* are proposed to assemble the glycosyl residues and catalyse their attachment to the thiopeptide core of the persiathiacins. The biosynthesis of these 6-deoxysugars is proposed to commence with the conversion of thymine diphosphate (TDP)-α-D-glucose **6** to TDP-4-keto-6-deoxy-α-D-glucose **7** catalysed by PerS2, which shows sequence similarity to TDP-glucose-4,6-dehydratases (Figure 5). At this point, the pathway appears to bifurcate. While PerS11, which is similar to TDP-hexose-4-ketoreductases, is proposed to catalyse the formation of TDP-6-deoxy-α-D-glucose **8** from TDP-4-keto-6-deoxy-α-D-glucose **7**, PerS10 and PerS1, which are homologues of TDP-4-keto-6-deoxy-D-glucose-2,3-dehydratases and TDP-4-keto-6-deoxy-D-glucose-3-ketoreductases, respectively, are hypothesised to convert **7** to TDP-4-keto-2,6-deoxy-α-D-glucose **9**. A further bifurcation then occurs. PerS11 catalyses the conversion of TDP-4-keto-2,6-deoxy-α-D-glucose **9** to TDP-D-olivose **10**, wherea**s** Per S12, which is similar to TDP-4-keto-2,6-deoxy-D-glucose-3-dehydratases converts **9** to TDP-4-keto-2,3,6-deoxy-α-D-glucose **10**. Finally, PerS11 catalyses the reduction of TDP-4-keto-2,3,6-deoxy-α-D-glucose **11** to TDP-D-amicetose **12**. This analysis is consistent with the assignment of sugars 2, 3 and 4 as D-amicetose, D-olivose, and 3, 4-di-O-methyl-6-deoxy-β-D-glucose, respectively, in persiathicin A **4**, and the substitution of D-amicetose by a second D-olivose residue in persiathiacin B **5**.

Philipimycin **3**, which bears the highest degree of structural similarity among known thiopeptide antibiotics to the persiathiacins, is proposed to be decorated with D-amecitose and two O-methylated L-rhamnose derivatives.^17^ Moreover, no 6-deoxysugar biosynthetic genes, beyond those hypothesised to be involved in the assembly of TDP-D-amicetose, TDP-D-olivose, and TDP-6-deoxy-β-D-glucose are present in the persiathiacin BGC (Figure 3 and Table S4). Because L-rhamnose is ubiquitously incorporated into bacterial cell surface carbohydrates,^44^ dedicated genes for TDP-L-rhamnose biosynthesis are invariably absent from rhamnosylated natural product BGCs. Taken together, these observations are consistent with the assignment of sugar 1 as an O-dimethylated L-rhamnose derivative. The deoxysugar residues contain a total of four methoxy groups (appended to C2 and C3 in sugar 1 and C3 and C4 in sugar 4), but only three putative O-methyltransferase-encoding genes (*perS3*, *perS5*, and *perS7*) are present in the persiathiacin BGC. It therefore appears that one of PerS3, PerS5 and PerS7 catalyses the O-methylation of two distinct hydroxyl groups in the biosynthesis of one of these sugars.

Four genes (*perS4*, *perS6*, *perS8*, and *perS9*) encode putative glycosyltransferases, each of which is hypothesised to append one glycosyl residue to the thiopeptide core. Glycosyltransferases are known to possess broad substrate tolerance,^45^ explaining why small amounts of persiathiacin B **5**, in which sugar 2 is D-olivose rather than D-amicetose, are produced in addition to persiathiacin A **4**.

**Figure 5.**
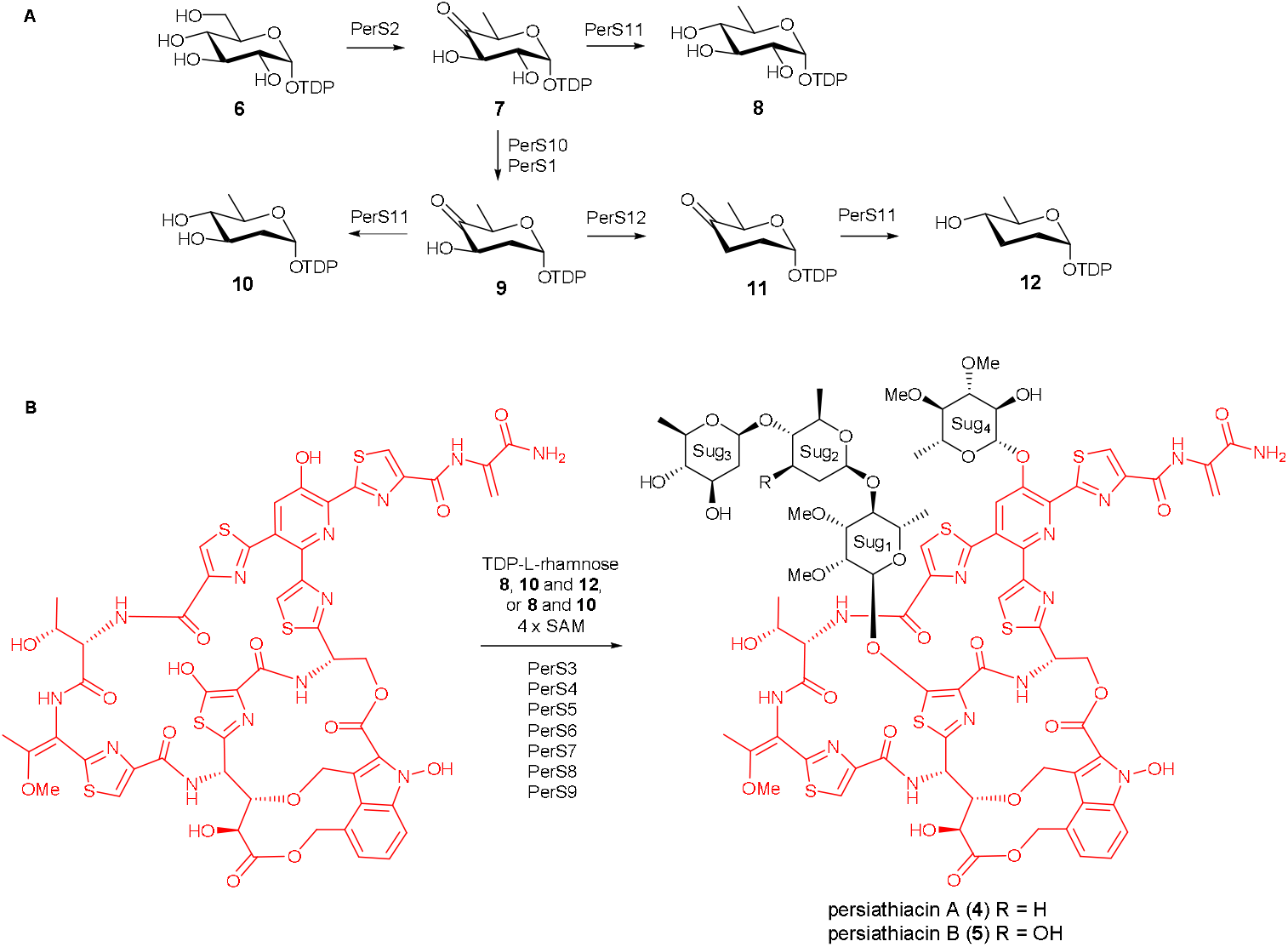
The enzymes encoded by *perS1-perS12* are proposed to be responsible for the biosynthesis and attachment of the 6-deoxysugars to the persiathiacin aglycone. (A) Proposed pathway for assembly of TDP-6-deoxy-α-D-glucose **8**, TDP-α-D-acemitose **10**, and TDP-α-D-acemitose **12**. (B) The glycosyltransferases encoded by *perS4*, *perS6*, *persS8* and *perS9* are proposed to decorate the persiathiacin core peptide with L-rhamnose, 6-deoxy-D-glucose, D-olivose and D-acemitose (persiathiacin A **4**), or L-rhamnose, 6-deoxy-D-glucose and D-olivose (persiathiacin B **5**). The methyl transferases encoded by *perS3*, *perS5* and *perS7* are hypothesized to O-methylate the L-rhamnose and 6-deoxy-D-glucose residues to produce the mature antibiotics. The timing of these transformations remains to be determined.

### PerX catalyses hydroxylation of nosiheptide *in vitro* and *in vivo*

Due to a lack of genetic tools for the *Actinokineospora* genus, we were unable to verify the involvement of the *Actinokineospora* sp. UTMC 2448 thiopeptide-like BGC in persiathiacin assembly via targeted disruption of one of the putative biosynthetic genes. Instead, we decided to investigate the proposed function of the putative CYP PerX as a novel type of thiopeptide-modifying enzyme. Recombinant His6-tagged PerX was overproduced in *E. coli* and purified using Nickel-affinity chromatography. The identity of the purified protein, including the presence of a haem prosthetic group, was confirmed by ESI-Q-TOF-MS analysis (Figure S16). The purified protein was incubated with commercially available nosiheptide **1,** spinach ferredoxin, spinach ferredoxin reductase, and NADPH at room temperature for 3 h. UHPLC-ESI-Q-TOF-MS analysis of the reaction mixture revealed a species with *m*/*z* = 1238.1493, corresponding to the [M+H]^+^ ion for a compound with the molecular formula C_51_H_43_N_13_O_13_S_6_ (calculated *m*/*z* = 1238.1500 for C_51_H_44_N_13_O_13_S_6_^+^) that was absent from a control reaction containing heat-inactivated enzyme. The molecular formula of this species is consistent with the insertion of an oxygen atom into nosiheptide (measured *m*/*z* = 1222.1542; calculated *m*/*z* = 1222.1551 for C_51_H_44_N_13_O_12_S_6_^+^) (Figure 6).

**Figure 6.**
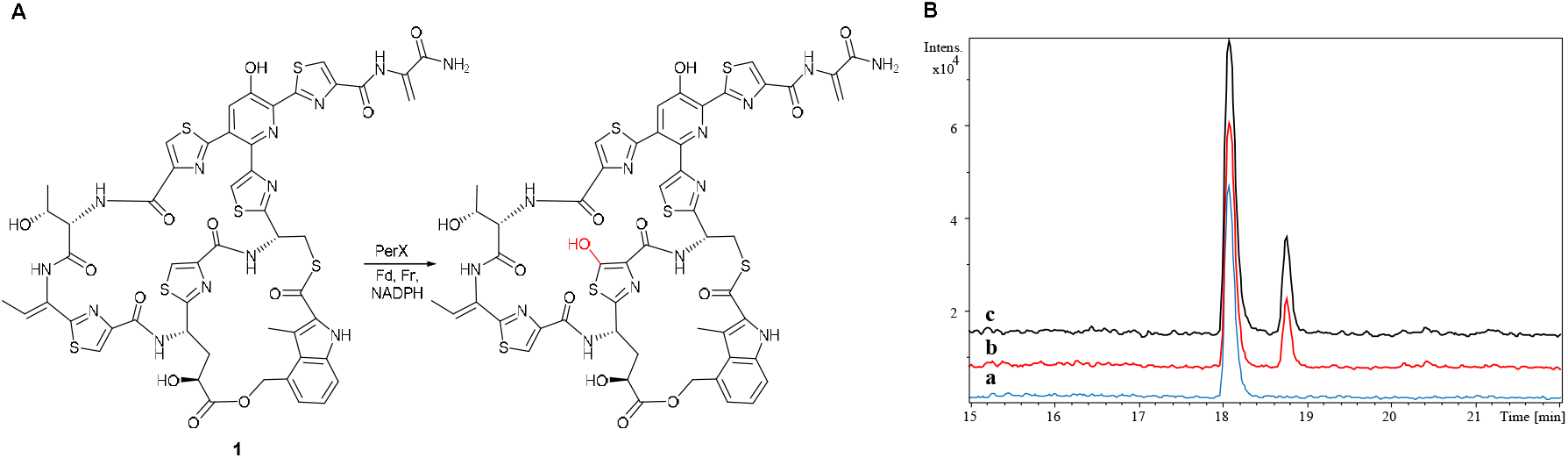
(A) Reaction catalysed by purified recombinant PerX with nosiheptide **1**, in the presence of spinach ferredoxin (Fd), spinach ferredoxin reductase (Fr) and NADPH. The proposed site of oxygen atom insertion, based on the assigned function of PerX in persiathiacin biosynthesis, is highlighted in red. (B) Extracted ion chromatograms at *m*/*z* = 1222.1551 and 1238.1500 corresponding to [M+H]^+^ for nosiheptide and its hydroxylated derivative, respectively, from UHPLC-ESI-Q-TOF-MS analyses of: culture extracts of *S. actuosus* ATCC25421 expressing *perX* under the control of the strong constitutive *ermE*\* promoter (top chromatogram); nosiheptide **1** after incubation for 3 hours with purified recombinant PerX, Fd, Fr and NADPH (middle chromatogram); and nosiheptide **1** after incubation for 3 hours with heat-denatured PerX, Fd, Fr and NADPH (bottom chromatogram).

In an attempt to obtain sufficient quantities of the oxygenated nosiheptide derivative for NMR spectroscopic analysis, *perX* was expressed under the control of the constitutive *ermE*\* promoter in the nosiheptide producer *Streptomyces actuosus* ATCC25421. Although the same nosiheptide derivative as that produced in the in vitro experiments was observed in UHPLC-ESI-Q-TOF-MS analyses of extracts from this strain (Figure 6), it was not possible to isolate sufficient quantities of this compound for full characterization by NMR spectroscopy.

### Biological activity and mode of action

Persiathiacin A was tested against the ESKAPE panel of pathogens by measuring minimum inhibitory concentrations (MICs).^46^ Persiathiacin A showed potent activity against MRSA (MIC of 0.025 μg/mL) and moderate activity against *Enterococcus faecium* (MIC of 32 μg/mL). The compound was inactive against all Gram-negative bacteria in the panel up to clinically relevant MIC cut offs. As nocathiacin has been reported to be active against drug-susceptible and resistant clinical strains of *M. tuberculosis*,^47, 48^ we evaluated the activity of persiathiacin A against several clinical isolates of *M. tuberculosis* using the resazurin microtiter assay.^49^ It was found to be active against all isolates tested, including the drug-susceptible strain H37Rv, four isoniazid-resistant strains, and two multi-drug resistant strains CHUV80059744 and CHUV80037024 - resistant to both isoniazid and rifampicin (Table 1). Persiathiacin A exhibited negligible toxicity toward the A2780 ovarian cancer cell line up to the maximum tested concentration of 400 μM (Figure S17).

**Table 1.** MIC values of persiathiacin A against several *M. tuberculosis* isolates.

| Strain | Mutation | Resistance | MIC (μg/mL) | | |
| --- | --- | --- | --- | --- | --- |
|  |  |  | Persiathiacin A | Rifampicin | Isoniazid |
| H37Rv | None | None | 3.6 | 0.0008 | 0.04 |
| HUGMB6726 | *inhA* | Isoniazid | 3.7 | 0.0008 | 2.2 |
| HUGMB3649 | *inhA* | Isoniazid | 3.9 | 0.001 | 0.08 |
| CHUV80045776 | *katG* | Isoniazid | 1.6 | 0.001 | 2.5 |
| HUGMI1020 | *katG* | Isoniazid | 1.7 | 0.002 | 1.3 |
| CHUV80059744 | *rpoB* and *katG* | Isoniazid, Rifampicin | 2.2 | 14 | >10 |
| CHUV80037024 | *rpoB*, *katG* and *inhA* | Isoniazid, Rifampicin | 3.1 | >10 | >10 |

The antibacterial activity of several thiopeptides is known to result from inhibition of ribosomal protein biosynthesis.^50–52^ We thus investigated the ability of persiathiacin A to inhibit the bacterial ribosome using an *E. coli* cell-free transcription/translation assay. Concentration-dependent inhibition of the ribosome by persiathiacin A was observed with an IC50 value of 1.8 ± 0.3 μM (Figure S18).

## Conclusions

Despite their potent antibacterial activity, the development of thiopeptides into clinically useful antibiotics has been prevented by their poor pharmacological properties, in particular their low aqueous solubility. Glycosylation is a widely used strategy for increasing the solubility of therapeutic peptides. However, most naturally occurring thiopeptides are either unglycosylated, or have a single sugar attached to the γ-hydroxyl group of the modified Glu residue, limiting opportunities to create novel glycosylated derivatives of thiopeptides via biosynthetic engineering. The sole exception, prior to our work, was philipimycin A, which has a trisaccharide appended to the central thiazole. Although philipimycin A is a rare example of thiopeptide that is active *in vivo*, nothing is known about its biosynthesis.

The discovery and biosynthetic elucidation of thiopeptides with novel glycosylation patterns, could provide a useful foundation for the creation of novel polyglycosylated thiopeptide derivatives with greater aqueous solubility and enhanced therapeutic potential. Our discovery in this work of the novel polyglycosylated thiopeptides, persiathiacins A and B from *Actinokineospora* sp. UTMC 2475 and *Actinokineospora* sp. UTMC 2448, and the gene cluster responsible for their biosynthesis, is therefore significant for several reasons. Firstly, the persiathiacins are the first examples of naturally occurring thiopeptides with a sugar appended to the hydroxpyridine. Glycosylation of the hydroxypyridine in nocathician has been reported to significantly improve aqueous solubility. The identification of the persiathiacin BGC opens the path for the development of biosynthetic engineering approaches to the creation of novel thiopeptide derivatives bearing glycosylated hydroxypyridines. Secondly, the identification of the persiathiacin BGC reveals the molecular mechanism for attachment of a trisaccharide to the central thiazole of thiopeptides. The incorporation of different sugars into the trisaccharides appended to the central thiazoles of phililipimycin A, persiathiacin A and persiathiacin B, suggests the glycosylation machinery is somewhat substrate tolerant. Thus, it seems plausible that biosynthetic engineering could be used to create a range of thiopeptide analogues with various mono, di and trisaccharides attached to the central thiazole. Thirdly, the observation that nocathiacin I, philipimycin A and persiathiacin A all display strong activity against *S. aureus* and *M. tuberculosis*, despite their diverse glycosylation patterns, indicates that creation and biological evaluation of novel glycosylated thiopeptide derivatives may be a fruitful strategy for circumventing the historical problems with this class of antibiotics that have prevented them from progressing into clinical application.

## Acknowledgements

The Bruker MaXis Impact UHPLC-ESI-Q-TOF-MS instrument used in this research was funded by the BBSRC (B/K002341/1 to G.L.C.). G.L.C. was the recipient of a Wolfson Research Merit Award from the Royal Society (WM130033). Y.D. was supported by a grant from the MRC (MR/N501839/1 to G.L.C.). We thank Ivan Prokes and Lijiang Song for assistance with recording NMR and UHPLC-ESI-Q-TOF-MS data, respectively. We thank Simone Schrader and Nicole Heyer for excellent technical assistance for PacBio genome sequencing.

