## Supplementary Information for "Discovery and biosynthesis of persiathiacins: Unusual polyglycosylated thiopeptides active against multi-drug resistant tuberculosis"

| **Contents** | **Page** |
| --- | --- |
| Materials and methods | S2-S9 |
| Figure S1. Structure of persiathiacin A, numbered as in Table S1. | S10 |
| Table S1.^1^H and ^13^C NMR chemical shifts of persiathiacin A in CDCl_3_-CD_3_OD (9:1). | S11-S13 |
| Figure S2. Structure of persiathiacin B, numbered as in Table S2. | S14 |
| Table S2.^1^H and ^13^C NMR chemical shifts of persiathiacin B in CDCl_3_-CD_3_OD (9:1). | S15-S17 |
| Figure S3. ^1^H NMR spectrum of persiathiacin A in CDCl_3_-CD_3_OD (9:1). | S18 |
| Figure S4. ^13^C NMR spectrum of persiathiacin A in CDCl_3_-CD_3_OD (9:1). | S19 |
| Figure S5. ^1^H-^1^H COSY spectrum of persiathiacin A in CDCl_3_-CD_3_OD (9:1). | S20 |
| Figure S6. ^1^H-^13^C HSQC spectrum of persiathiacin A in CDCl_3_-CD_3_OD (9:1). | S21 |
| Figure S7. ^1^H-^13^C HMBC spectrum of persiathiacin A in CDCl_3_-CD_3_OD (9:1). | S22 |
| Figure S8. ^1^H-^1^H ROSY spectrum of persiathiacin A in CDCl_3_-CD_3_OD (9:1). | S23 |
| Figure S9. ^1^H NMR spectrum of persiathiacin B in CDCl_3_-CD_3_OD (9:1). | S24 |
| Figure S10. ^13^C NMR spectrum of persiathiacin B in CDCl_3_-CD_3_OD (9:1). | S25 |
| Figure S11. ^1^H-^1^H COSY spectrum of persiathiacin B in CDCl_3_-CD_3_OD (9:1). | S26 |
| Figure S12. ^1^H-^13^C HSQC spectrum of persiathiacin B in CDCl_3_-CD_3_OD (9:1). | S27 |
| Figure S13. ^1^H-^13^C HMBC spectrum of persiathiacin B in CDCl_3_-CD_3_OD (9:1). | S28 |
| Figure S14. ^1^H-^1^H ROSY spectrum of persiathiacin B in CDCl_3_-CD_3_OD (9:1). | S29 |
| Table S3: Predicted secondary metabolite biosynthetic gene clusters in the genome of *Actinokineospora* sp. UTMC 2448. | S30 |
| Table S4: Genes/proteins in the persiathiacin biosynthetic gene cluster and their similarity to homologous proteins. | S31- S32 |
| Figure S15. Similarities and differences in the activation and attachment of MIA to the nosiheptide, persiathiacin, and nocathiacin processed core peptides. | S33 |
| Figure S16. Mass spectrum of PerX | S34 |
| Figure S17. Percentage of A2780 ovarian cancer cells surviving after exposure to increasing concentrations of persiathiacin A (10 to 400 μM) relative to an untreated control. | S35 |
| Figure S18. Concentration-dependent inhibition of the *E. coli* ribosome by persiathiacin A. | S35 |

**Materials and Methods**

**General Experimental Procedures:** Optical rotations were measured on an Optical Activity Ltd AA-1000 millidegree auto-ranging polarimeter (589 nm). Specific rotations are given in units of 10^-1^ deg cm^2^ g^-1^. UV spectra were acquired on a Perkin Elmer Lambda 35 UV/Vis spectrophotometer. IR spectra were recorded on an Alpha Bruker Platinum ATR single reflection diamond ATR module. UHPLC-ESI-Q-TOF-MS analyses were performed using a Dionex UltiMate 3000 UHPLC connected to a Zorbax Eclipse Plus C18 column (100 × 2.1 mm, 1.8 μm) coupled to a Bruker MaXis IMPACT mass spectrometer. Mobile phases consisted of water (A) and acetonitrile (B), each supplemented with 0.1% formic acid. A gradient of 5 to 100% B over 30 minutes was employed at a flow rate of 0.2 mL/min. The mass spectrometer was operated in positive ion mode with a scan range of 50-3000 *m/z*. Calibration was performed with 1 mM sodium formate through a loop injection of 20 μL at the start of each run. Persiathiacin A and B were dissolved in a mixture of CDCl_3_-CD_3_OD (9:1) for NMR spectroscopic analyses. NMR spectra were recorded on Bruker 500 or 700 MHz spectrometers equipped with DUI and TCl cryoprobes, respectively, at 25 °C. The ^1^H and ^13^C NMR chemical shifts were referenced to the solvent peaks at *δ*_H_ 7.26 and *δ*_C_ 77.16 for CDCl_3_. All HPLC and LC-MS experiments were performed with the MeCN-H_2_O gradient solvent system. Millipore Milli-Q H_2_O and HPLC grade solvents were used for chromatography.

**Strain Isolation and Identification:** The actinobacteria *Actinokineospora* sp. UTMC 2475 and *Actinokineospora* sp. UTMC 2448 were isolated from mud samples collected from Bushehr and Chabahar, Iran. The samples were dried at 50 °C, ground to a powder and passed through a 2 mm sieve. Strains were isolated on solid Reasoner’s 2A (R2A) medium^1^ after 3 weeks of incubation at 28 °C. Solid ISP2 medium was then used to purify the strains. Purified strains were preserved in 30% glycerol at -70 °C. To identify the strains, 16S rRNA genes were amplified using a set of universal primers (27F, 1100F, 1100R, 1525R). Amplified DNA obtained from the reactions was purified using a PCR purification kit (Roti®-Prep PCR Purification). The 16S rRNA gene sequence of the strains was BLASTed against the GenBank and EzTaxon databases.^2^

**Production, Extraction, and HPLC Purification of Persiathiacins A and B:** *Actinokineospora* sp. UTMC 2448 was grown on solid ISP2 medium (4 g/L glucose, 4 g/L yeast extract, 10 g/L malt extract, 2 g/L CaCO_3_, 15 g/L Bacto agar) for 7 days at 30 °C. The agar cultures were chopped and extracted with EtOAc. The extract was dried on a rotary evaporator and pre-adsorbed to C18-bonded silica, and then packed into a stainless steel HPLC guard cartridge (10 × 30 mm) attached to a semi-preparative reverse-phase C18 Betasil column (21.2 mm × 150 mm). The column was eluted with 5% acetonitrile for 5 min, then a linear gradient from 5 to 100% acetonitrile was applied over 45 min, and the columns was eluted for an additional 10 min with 100% acetonitrile. The flow rate was 9 mL/min. Sixty fractions were collected in 1 min increments over 60 min. Pure persiathiacin A was obtained in fraction 35. Persiathiacin B was purified from a mixture of persiathiacin A and B in fraction 34 using a reverse-phase C18 Betasil column (21.2 mm × 150 mm). Isocratic elution with 40% acetonitrile for 5 min followed by a linear gradient to 65% over 45 min was used to achieve separation of persiathiacin B (fraction 26) from persiathiacin A (fraction 28).

*Persiathiacin A*: amorphous solid (19 mg); [α]_D_^26^ +86 (*c* 0.05, CHCl_3_:MeOH (9:1)); UV (CHCl_3_:MeOH (9:1)) λ_max_ (log ε) 231 (5.85), 278 (5.65), 353 (5.25) nm; IR *ν*_max_ 3392, 2980, 1725, 1667, 1534, 1250, 1205, 1121, 1065, 751 cm^−1^; ^1^H NMR (500 MHz, CDCl_3_:CD_3_OD (9:1)) and ^13^C NMR (125 MHz, CDCl_3_:CD_3_OD (9:1)), see Table S1; HRESIMS *m*/*z* 1874.4638 [M + H]^+^ (calcd for C_80_H_92_N_13_O_30_S_5_, 1874.4671).

*Persiathiacin B*: amorphous solid (1.8 mg); [α]_D_^26^ +86 (*c* 0.05, CHCl_3_:MeOH (9:1)); UV (CHCl_3_:MeOH (9:1)) λ_max_ (log ε) 232 (5.86), 275 (5.61), 353 (5.21) nm; IR *ν*_max_ 3402, 2970, 1667, 1533, 1250, 1205, 1121, 1067, 1037, 751 cm^−1^; ^1^H NMR (700 MHz, CDCl_3_:CD_3_OD (9:1)) and ^13^C NMR (175 MHz, CDCl_3_:CD_3_OD (9:1)), see Table S1; HRESIMS *m*/*z* 1890.4611 [M + H]^+^ (calcd for C_80_H_92_N13O_31_S_5_, 1890.4620).

**PacBio library preparation and sequencing:** Genomic DNA was extracted from *Actinokineospora* sp. UTMC2448. A SMRTbell™ template library was prepared according to the manufacturer’s instructions (Pacific Biosciences, Menlo Park, CA, USA), following the Procedure & Checklist – Greater Than 10 kb Template Preparation. Briefly, for preparation of 15 kb libraries, 8 µg genomic DNA was sheared using g-tubes™ (Covaris, Woburn, MA, USA) according to the manufacturer´s instructions. DNA was end-repaired and ligated overnight to hairpin adapters by applying components from the DNA/Polymerase Binding Kit P6 (Pacific Biosciences, Menlo Park, CA, USA). Reactions were carried out according to the instructions of the manufacturer. BluePippin™ Size-Selection to greater than 4 kb was performed according to the manufacturer´s instructions (Sage Science, Beverly, MA, USA). Conditions for annealing of sequencing primers and binding of polymerase to purified SMRTbell™ template were assessed with the Calculator in RS Remote (Pacific Biosciences, Menlo Park, CA, USA). SMRT sequencing was carried out on the PacBio *RSII* (Pacific Biosciences, Menlo Park, CA, USA), taking one 240-minute movie for one SMRT cell using the P6 Chemistry. Sequencing resulted in 76,562 post-filtered reads with a mean read length of 10,180 bp.

**Genome assembly, error correction, and annotation:** SMRT cell data was assembled using the “RS_HGAP_Assembly.3” protocol included in SMRT Portal version 2.3.0 using default parameters. The assembly resulted in a single circular chromosome. Error correction was performed by a mapping of 7 million paired-end Illumina reads of 2 x 100 bp onto the genome using BWA (PMID 19451168)^3^ with subsequent variant and consensus calling using VarScan (PMID 22300766).^4^ A consensus concordance of QV60 could be confirmed for the genome. Finally, annotation was carried out using Prokka 1.8 (PMID 24642063).^5^ Prediction of secondary metabolite biosynthetic gene clusters was made using antiSMASH v3.0 (Table S3). The putative persiathiacin gene cluster was subjected to detailed manual annotation via comparative sequence analysis (Table S4).

**Overproduction and Purification of PerX:** The gene encoding PerX was PCR-amplified from *Actinokineospora sp.* UTMC 2448 gDNA using Phusion DNA polymerase (NEB) and primers 5′-GTGCCGCGCGGCAGCCATATGCTTCCCGAGCCGTACACCCCCGAGTTCT-3′ and 5′-TCGACGGAGCTCGAATTCTCATCGCGTCACCCGCAGCTCGGCCA-3′ (regions complementary to the gene sequence underlined). The linear pET28a (NEB) vector backbone was PCR-amplified with primers 5′-TGAGAATTCGAGCTCCGTCGACAAGCTTG-3′ and 5′-CATATGGCTGCCGCGCGGCAC-3′. PCR products were separated on a 1% agarose gel and bands were excised and purified with a GeneJET Gel Extraction Kit (Thermo Scientific). Cloning of the pure insert into the *Nde*I/*Eco*RI restriction sites of the linear pET28a vector was accomplished by Gibson assembly following the manufacturer’s instructions (NEB). The resulting vector was used to transform *E. coli* TOP10 cells (Invitrogen) and plated on LB agar containing kanamycin (50 μg/mL). Colonies were picked and grown overnight in liquid LB medium. Plasmids were isolated from the culture using a GeneJET Plasmid Miniprep Kit (Thermo Scientific) and inserts were sequenced to verify their integrity. The correct pET28a plasmid containing *perX* was used to transform *E. coli* BL21(DE3) cells. A single colony was used to inoculate liquid LB medium (10 mL) containing kanamycin (50 μg/mL), which was incubated overnight at 37 °C and 180 rpm; this was then used to further inoculate liquid LB medium (1 L) containing kanamycin (50 μg/mL). The resulting culture was incubated at 37 °C and 180 rpm until OD_595nm_ reached 0.6, then IPTG (0.5 mM) was added, and expression was continued overnight at 15 °C and 180 rpm. The cells were harvested by centrifugation (5,000 rcf, 20 min, 4 °C) and resuspended in buffer (30 mM HEPES, 500 mM NaCl, 10% glycerol, pH 7.5) at 20 mL/L of growth medium, then lysed using sonication (Vibra-Cell Ultrasonic Liquid Processor; Sonics & Materials, Inc.). The lysate was centrifuged (30,000 rcf, 60 min, 4 °C) and the resulting supernatant was passed through a 0.45 µm filter (Sartorius). An ÄKTA pure FPLC (GE Healthcare) was used to purify PerX as follows. The supernatant was loaded onto a 1-mL HisTrap HP column (GE Healthcare), which had been equilibrated with resuspension buffer (30 mM HEPES, 500 mM NaCl, 10% glycerol, pH 7.5). Proteins were eluted in a stepwise manner using increasing concentrations of imidazole (0 to 150 mM) in resuspension buffer. The presence of the protein of interest in the elution fractions was confirmed by SDS-PAGE. Fractions containing the pure protein were pooled and concentrated to ~100 μM using a 50 kDa MWCO Vivaspin centrifugal concentrator (Sartorius). Aliquots of 50 μL were snap-frozen in liquid N_2_ and stored at -80 °C until further use.

**Hydroxylation of Nosiheptide by PerX:** A 200 μL reaction mixture containing nosiheptide (100 μM), spinach ferredoxin-NADP^+^ reductase (0.1 U/mL), spinach ferredoxin (50 μg/mL), NADPH (1 mM), and PerX (10 μM) in Tris-HCl (25 mM, pH 8) was incubated at room temperature for 3 hours. The reaction was terminated by adding 200 μL of methanol, and after separating the precipitate by centrifugation (16,000 rcf, 10 min) the supernatant was analysed by UHPLC-ESI-Q-TOF-HRMS. For the negative control, PerX was inactivated by boiling at 100 °C for 15 min.

**Expression of *per*X in the Nosiheptide-Producing Strain *Streptomyces actuosus*:** *perX* was amplified from *Actinokineospora sp.* UTMC 2448 gDNA using Phusion DNA polymerase (NEB) and primers 5′-CAGCATATGGTGCTTCCCGAGCCGTAC-3′ and 5′-GACGAATTCTCATCGCGTCACCCGC-3′. The PCR product was digested with *Nde*I and *Eco*RI and cloned into the corresponding sites of pIB139 under the control of the *ermE** constitutive promoter. The integrity of the construct was confirmed by sequencing and the resulting plasmid was used to transform *E. coli* ET12567/pUZ8002 cells by electroporation. A mixture of apramycin (50 μg/mL), kanamycin (50 μg/mL) and chloramphenicol (35 μg/mL) was used for selection on LB agar. The pIB139 vector containing *per*X was then introduced by conjugation into *Streptomyces actuosus* ATCC25421. The overnight culture was plated on SFM agar medium and overlaid for with 1 ml of antibiotic solution mixture containing apramycin (50 μg/mL) and nalidixic acid (25 μg/mL). After 3 days, four colonies were picked and spread separately onto SFM agar medium containing apramycin (50 μg/mL) and nalidixic acid (25 μg/mL) and then further sub-cultured on five plates for to produce spores. Spores from the resulting stocks were cultured in liquid medium containing corn steep liquor (10 g/L), soy flour (20 g/L), yeast extract (3 g/L), NaCl (4 g/L), KNO_3_ (0.2 g/L), CaCO_3_ (4 g/L), pH 7.0. Production of the hydroxylated nosiheptide derivative was confirmed by UHPLC-ESI-Q-TOF-MS analysis.

**MIC assays against *M. tuberculosis*:** Dimethyl sulfoxide (DMSO), glycerol, isoniazid, resazurin sodium salt, and rifampicin were purchased from Sigma-Aldrich (USA). Middlebrook 7H9 was purchased from Difco (USA) and albumin dextrose catalase (ADC) from Chemie Brunschwig AG (Switzerland). The *M. tuberculosis* reference strain H37Rv was obtained from Institut Pasteur, Paris, and clinical specimens from patients were obtained from the Lausanne University Hospital (CHUV) and Geneva University Hospital (HUG).

The resazurin reduction microplate assay (REMA) was performed as described previously.^6^ Two-fold serial dilutions of each test compound were prepared in 96-well plates from 10 mg/mL stocks in DMSO. Frozen aliquots of replicating tubercule bacilli (reference strains and clinical isolates) were thawed and diluted to an OD_600_ of 0.0001 (3 · 10^4^ cells/mL) and added to the plates to obtain a total volume of 100 μL. Plates were incubated for 6 days at 37 °C before adding resazurin (0.025% [w/v] to 1/10 of well volume). After overnight incubation, fluorescence of the resazurin metabolite resorufin was determined by excitation at 560 nm and emission at 590 nm, as measured by a TECAN infinite M200 microplate reader. The MIC was defined visually as the lowest concentration to prevent resazurin turnover from blue to pink and was confirmed by the level of measured fluorescence. MIC values were calculated using GraphPad Prism version 7.0 (GraphPad Software, Inc., La Jolla, CA, USA). The experiment was performed twice, and all the compounds were tested in triplicate (total of six replicates).

**Ribosome inhibition assay:** The ability of persiathiacin to inhibit ribosomal protein synthesis was assessed in an *E. coli* cell-free transcription/translation assay in which production of firefly luciferase was monitored in the presence of increasing concentrations of persiathiacin. The assay was carried out in triplicate as previously described.^7^ IC_50_ values were determined by least-squares regression analysis as implemented in the GraphPad Prism software package.

**Cytotoxicity assays:** Evaluation of the cytotoxicity of persiathiacin A was carried out using A2780 ovarian cancer cells, which were obtained from the European Collection of Cell Cultures. Cells were grown as adherent monolayers using Roswell Park Memorial Institute medium (RPMI 1640) supplemented with 10% v/v of foetal calf serum, 1% v/v of 2 mM glutamine and 1% v/v penicillin/streptomycin using a 5% CO_2_ humidified atmosphere. Cultures were regularly passaged when achieving 70-80% confluence. For these experiments, cells were seeded in a 96 well plate at a density of 5000 cells/well and they were allowed to attach for 48h in persiathiacin-free medium. Various concentrations of persiathiacin were added in concentrations up to 400 µM. Working solutions were obtained by dilution with cell culture medium from a 5% v/v DMSO:RPMI stock. After 24 h of drug exposure, cells were washed, and fresh medium was replenished to allow for 72 h of recovery time. Cell viability was assessed using the MTT assay. Formazan absorbance at 570 nm was recorded in a FLUOstar Omega microplate reader. In all cases, reported values were obtained as duplicates of triplicates in independent experiments with their associated standard deviations.

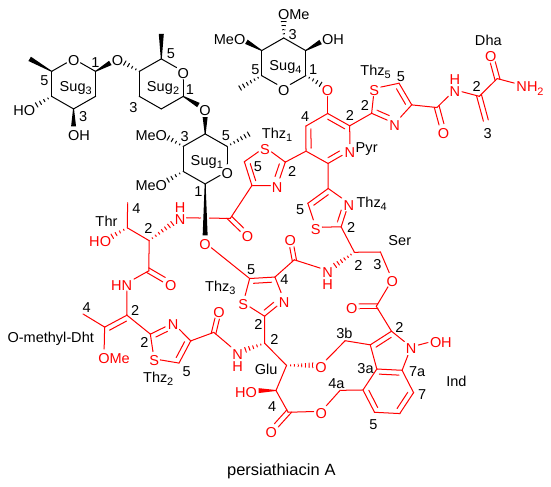

**Figure S1.** Structure of persiathiacin A, numbered as in Table S1.

**Table S1.** ^1^H (500 MHz) and ^13^C (125 MHz) and HMBC NMR data for persiathiacin A in CDCl_3_-CD_3_OD (9:1).

| Position | | *δ*_C_ | *δ*_H_ | HMBC | ROESY |
| --- | --- | --- | --- | --- | --- |
| Dha | C=O | 166.5 |  |  |  |
|  | C2 | 133.5 |  |  |  |
|  | C3 | 104.8 | H3a: 5.5, brs  H3b: 6.5, brs | Dha-(C=O)  Dha-(C2, C=O) | Dha-(H3b)  Dha-(H3a) |
|  | NH |  | 9.98, s | Dha-(C3, C=O), Thz5-(C4, C=O) |  |
| Thz5 | C2 | 166.9 |  |  |  |
|  | C4 | 151.2 |  |  |  |
|  | C5 | 127.2 | 8.15, s | Thz5-(C2, C4, C=O) |  |
|  | C=O | 160.0 |  |  |  |
| Pyr | C2 | 139.8 |  |  |  |
|  | C3 | 149.0 |  |  |  |
|  | C4 | 126.7 | 7.77, s | Pyr-(C2, C3, C6), Thz-(C2) | Sug4-(H1) |
|  | C5 | 128.3 |  |  |  |
|  | C6 | 144.8 |  |  |  |
| Thz1 | C2 | 165.5 |  |  |  |
|  | C4 | 149.1 |  |  |  |
|  | C5 | 125.2 | 8.25, s | Thz1-(C2, C4, C=O) | Thr-(H4) |
|  | C=O | 161.8 |  |  |  |
| Thr | NH |  | 7.87, brs | Thz1-(C=O) |  |
|  | C=O | 167.5 |  |  |  |
|  | C2 | 56.2 | 4.21, dd, *J* = 4.0, 5.5 | Thr-(C3, C4, C=O), Thz1-(C=O) | Thr-(H3)  Dht-(NH) |
|  | C3 | 64.7 | 2.88, m | Thr-(C=O) | Thr-(H2) |
|  | C4 | 17.7 | 1.34, d, *J* = 6.0 | Thr-(C2, C3) | Thz1-(H5) |
| O-methyl-Dht | NH |  | 8.31, s | Dht-(C3), Thr-(C=O) | Thr-(H2) |
|  | C2 | 110.7 |  |  |  |
|  | C3 | 158.7 |  |  |  |
|  | C4 | 13.7 | 1.86, s | Dht-(C2, C3), Thz2-(C2) | Dht-(NH, OCH_3_), Thr-(H2) |
|  | OMe | 55.8 | 3.78, s | Dht-(C3) | Dht-(H4) |
| Thz2 | C2 | 161.8 |  |  |  |
|  | C4 | 146.1 |  |  |  |
|  | C5 | 124.3 | 7.96, s | Thz2-(C2, C4, C=O), Dht-(C2) |  |
|  | C=O | 161.7 |  |  |  |
| Glu | NH |  | 8.24, brs | Glu-(C2, C3), Thz2-(C=O) | Glu-(H2, H4) |
|  | C2 | 48.7 | 5.80, d, *J* = 10.0 | Glu-(C3, C4), Thz3-(C2) | Glu-(NH, H3, H4) |
|  | C3 | 82.9 | 3.65, m | Glu-(C4, C=O), Thz3-(C2), Ind (C3b) | Glu-(H2), Ind-(H3ba/b) |
|  | C4 | 67.7 | 4.09, d, *J* = 10.0 | Glu-(C2, C3, C=O) | Glu-(NH, H2, H3) |
|  | C=O | 174.5 |  |  |  |
| Thz3 | C2 | 154.8 |  |  |  |
|  | C4 | 130.9 |  |  |  |
|  | C5 | 160.3 |  |  |  |
|  | C=O | 160.3 |  |  |  |
| Ser | NH |  | 7.89, d, *J* = 11.0 | Ser-(C2, C3), Thz3-(C4, C=O) | Ser-(H2, H3b) |
|  | C2 | 48.6 | 5.62, dd, *J* = 6.0, 10.5 | Ser-(C3), Thz3-(C=O), Thz4-(C2) | Ser-(NH, H3a/b) |
|  | C3 | 64.5 | H3a: 4.36, d, *J* = 11.5  H3b: 5.21, dd, *J* = 6.0, 11.5 | Ser-(C2), Thz4-(C2), Ind-(C=O)  Ser-(C2), Thz4-(C2), Ind-(C=O) | Ser-(H2, H3b)  Ser-(NH, H2, H3a) |
| Thz4 | C2 | 169.8 |  |  |  |
|  | C4 | 154.7 |  |  |  |
|  | C5 | 121.2 | 7.77, s | Thz4-(C2, C4), Pyr-(C6) |  |
| Ind | C=O | 161.4 |  |  |  |
|  | C2 | 127.0 |  |  |  |
|  | C3 | 109.8 |  |  |  |
|  | C3a | 119.5 |  |  |  |
|  | C3b | 65.9 | H3ba: 4.17, d, *J* = 10.5  H3bb: 4.91, d, *J* = 10.5 | Ind-(C2, C3, C3a), Glu-(C3)  Ind-(C2, C3, C3a) | Glu-(H3), Ind-(H3bb, H4ab)  Glu-(H3), Ind-(H3ba), Ser-(NH) |
|  | C4 | 127.8 |  |  |  |
|  | C4a | 68.4 | H4aa: 4.95, d, *J* = 12.5  H4ab: 5.92, d, *J* = 12.5 | Ind-(C3a, C4, C5), Glu-(C=O)  Ind-(C3a, C4, C5), Glu-(C=O) | Ind-(H3ba, H4ab, H5)  Ind-(H3ba, H4aa, H5) |
|  | C5 | 123.5 | 7.10, d, *J* = 7.0 | Ind-(C3a, C4a, C6, C7) | Ind-(H4aa, H4ab, H6) |
|  | C6 | 125.1 | 7.35, dd, *J* = 7.0, 8.5 | Ind-(C4, C5, C7, C7a) | Ind-(H5, H7) |
|  | C7 | 112.1 | 7.74, d, *J* = 8.5 | Ind-(C3a, C5) | Ind-(H6) |
|  | C7a | 135.3 |  |  |  |
|  | N-OH |  | 10.46, s | Ind-(C2) |  |
| Sug1 | C1 | 101.9 | 5.41, brs | Thz3-(C5), Sug1-(C3, C5) | Sug1-(H2, H4, C2-OCH_3_) |
|  | C2 | 76.0 | 4.00, m | Sug1-(C1, C3, C4, C2-OMe) | Sug1-(H1, H3, C2-OCH_3_) |
|  | C2-OMe | 59.4 | 3.48, s | Sug1-(C2) | Sug1-(H1) |
|  | C3 | 80.1 | 3.60, m | Sug1-(C1, C5, C3-OMe) | Sug1-(H2, C3-OCH_3_) |
|  | C3-OMe | 57.9 | 3.42, s | Sug1-(C3) | Sug1-(H2, H4) |
|  | C4 | 77.0 | 3.64, m | Sug1-(C2, C5, C6), Sug2-(C1) | Sug2-(H1) |
|  | C5 | 70.1 | 3.63, m | Sug1-(C1, C3, C4, C6) | Sug1-(H2, H6) |
|  | C6 | 17.7 | 1.31, d, *J* = 5.5 | Sug1-(C4, C5) | Sug1-(H4) |
| Sug2 | C1 | 102.2 | 4.61, d, *J* = 9.0 | Sug1-(C4), Sug2-(C2, C5) | Sug1-(H4), Sug2-(H2b, H3a, H5) |
|  | C2 | 30.8 | H2a: 1.39, m  H2b: 1.79, m | Sug2-(C1, C3, C4) | Sug2-(H1, H2b H3b, H4)  Sug2-(H1, H2a H3b) |
|  | C3 | 29.9 | H3a: 1.40, m  H3b: 2.09, m | Sug2-(C1, C4) | Sug2-(H2b, H4) |
|  | C4 | 80.6 | 3.08, m | Sug2-(C5, C6), Sug3-(C1) | Sug3-(H1), Sug2-(H3a/b, H6) |
|  | C5 | 74.1 | 3.26, m | Sug2-(C1, C3, C6) | Sug2-(H1, H3b, H6) |
|  | C6 | 18.0 | 1.12, d, *J* = 6.0 | Sug2-(C4, C5) | Sug2-(H4, H5), Sug3-(H1) |
| Sug3 | C1 | 101.1 | 4.43, d, *J* = 9.5 | Sug2-(C4), Sug3-(C2, C3, C5) | Sug2-(H4, H6), Sug3-(H2a, H3, H5) |
|  | C2 | 39.0 | H2a: 1.46, m  H2b: 2.06, m | Sug3-(C1, C3, C4) | Sug3-(H2b, H4)  Sug3-(H1, H2a, H3) |
|  | C3 | 71.3 | 3.42, *ol*^*^ | Sug3-(C1, C5) | Sug3-(H1, H2b) |
|  | C4 | 77.0 | 2.90, m | Sug3-(C2, C5, C6) | Sug3-(H2a, H3, H6) |
|  | C5 | 71.8 | 3.14, m | Sug3-(C1, C4, C6) | Sug3-(H1, H3 H6) |
|  | C6 | 17.7 | 1.20, d, *J* = 6.0 | Sug3-(C4, C5) | Sug3-(H4, H5) |
| Sug4 | C1 | 100.4 | 5.18, d, *J* = 7.5 | Pyr-(C3), Sug4-(C3, C5) | Pyr-(H4), Sug4-(H2, H3, H5) |
|  | C2 | 69.5 | 4.14, dd, *J* = 7.5, 10.0 | Sug4-(C1, C3) | Sug4-(H1, C3-OCH_3_) |
|  | C3 | 84.2 | 3.17, *ol* | Sug4-(C2, C4, C3-OMe) | Sug4-(H1, H5) |
|  | C3-OMe | 58.1 | 3.45, s | Sug4-(C3) |  |
|  | C4 | 77.6 | 3.42, *ol* | Sug4-(C2, C4-OMe) | Sug4-(H6) |
|  | C4-OMe | 61.8 | 3.51, s | Sug4-(C4) | Sug4-(H6) |
|  | C5 | 71.3 | 3.76, m | Sug4-(C1, C3, C4, C6) | Sug4-(H1, H4, H6) |
|  | C6 | 16.7 | 1.30, d, *J* = 6.0 | Sug4-(C4, C5) |  |

^* overlapped^

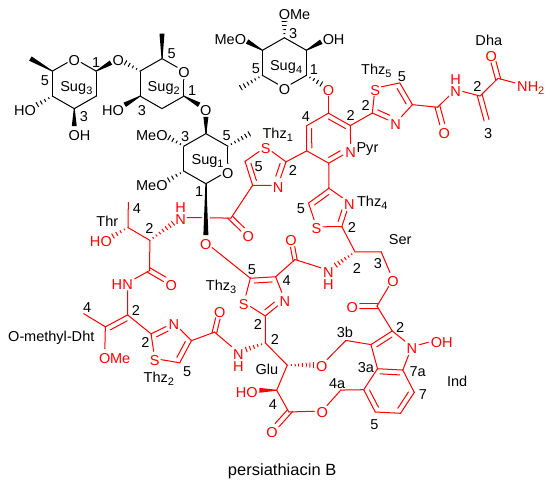

**Figure S2.** Structure of persiathiacin B, numbered as in Table S2.

**Table S2.** ^1^H (700 MHz) and ^13^C (176 MHz) and HMBC NMR data of persiathiacin B in CDCl_3_-CD_3_OD (9:1).

| Position | | | *δ*_C_ | | *δ*_H_ | | HMBC | ROESY |
| --- | --- | --- | --- | --- | --- | --- | --- | --- |
| Dha | C=O | 166.7 | |  | |  | |  |
|  | C2 | 133.5 | |  | |  | |  |
|  | C3 | 105.1 | | H3a: 5.56, brs  H3b: 6.51, brs | | Dha-(C=O)  Dha-(C2, C=O) | | Dha-(H3b)  Dha-(H3a) |
|  | NH |  | | 9.98, s | | Deala-(C3, C=O), Thz5-(C4, C=O) | |  |
| Thz5 | C2 | 167.1 | |  | |  | |  |
|  | C4 | 151.2 | |  | |  | |  |
|  | C5 | 127.4 | | 8.21, s | | Thz5-(C2, C4, C=O) | |  |
|  | C=O | 160.1 | |  | |  | |  |
| Pyr | C2 | 139.9 | |  | |  | |  |
|  | C3 | 149.1 | |  | |  | |  |
|  | C4 | 126.9 | | 7.79, s | | Pyr-(C2, C3, C6), Thz-(C2) | | Sug4-(H1) |
|  | C5 | 128.4 | |  | |  | |  |
|  | C6 | 145.0 | |  | |  | |  |
| Thz1 | C2 | 165.5 | |  | |  | |  |
|  | C4 | 149.1 | |  | |  | |  |
|  | C5 | 125.2 | | 8.27, s | | Thz1-(C2, C4, C=O) | | Thr-(H4) |
|  | C=O | 161.9 | |  | |  | |  |
| Thr | NH |  | | 7.94, brs | | Thz1-(C=O) | |  |
|  | C=O | 167.5 | |  | |  | |  |
|  | C2 | 56.2 | | 4.26, t, *J* = 4.0 | | Thr-(C3, C4, C=O), Thz1-(C=O) | | Thr-(H3)  Dht-(NH) |
|  | C3 | 64.8 | | 2.90, m | | Thr-(C=O) | | Thr-(H2) |
|  | C4 | 17.6 | | 1.37, d, *J* = 6.0 | | Thr-(C2, C3) | | Thz1-(H5) |
| O-methyl-Dht | NH |  | | 8.35, s | | Dht-(C3), Thr-(C=O) | | Thr-(H2) |
|  | C2 | 110.7 | |  | |  | |  |
|  | C3 | 158.8 | |  | |  | |  |
|  | C4 | 13.7 | | 1.89, s | | Dht-(C2, C3), Thz2-(C2) | | Dht-(NH, OCH_3_), Thr-(H2) |
|  | OMe | 55.9 | | 3.81, s | | Dht-(C3) | | Dht-(H4) |
| Thz2 | C2 | 161.8 | |  | |  | |  |
|  | C4 | 146.1 | |  | |  | |  |
|  | C5 | 124.4 | | 7.99, s | | Thz2-(C2, C4, C=O), Dht-(C2) | |  |
|  | C=O | 161.8 | |  | |  | |  |
| Glu | NH |  | | 8.27, brs | | Glu-(C2, C3), Thz2-(C=O) | | Glu-(NH, H2, H4) |
|  | C2 | 48.7 | | 5.82, dd, *J* = 1.5, 10.0 | | Glu-(C3, C4), Thz3-(C2) | | Glu-(H3, H4) |
|  | C3 | 83.0 | | 3.66, m | | Glu-(C4, C=O), Thz3-(C2), Ind (C3b) | | Glu-(H2), Ind-(H3ba/b) |
|  | C4 | 67.8 | | 4.12, d, *J* = 10.0 | | Glu-(C2, C3, C=O) | | Glu-(NH, H2, H3) |
|  | C=O | 174.5 | |  | |  | |  |
| Thz3 | C2 | 154.7 | |  | |  | |  |
|  | C4 | 131.0 | |  | |  | |  |
|  | C5 | 160.3 | |  | |  | |  |
|  | C=O | 160.3 | |  | |  | |  |
| Ser | NH |  | | 7.91, d, *J* = 11.0 | | Ser-(C2, C3), Thz3-(C4, C=O) | | Ser-(H2, H3b) |
|  | C2 | 48.6 | | 5.66, dd, *J* = 6.0, 11.0 | | Ser-(C3), Thz3-(C=O), Thz4-(C2) | | Ser-(NH, H3a/b) |
|  | C3 | 64.4 | | H3a: 4.40, d, *J* = 11.5  H3b: 5.25, dd, *J* = 6.0, 11.5 | | Ser-(C2), Thz4-(C2), Ind-(C=O)  Ser-(C2), Thz4-(C2), Ind-(C=O) | | Ser-(H2, H3b)  Ser-(NH, H2, H3a) |
| Thz4 | C2 | 169.9 | |  | |  | |  |
|  | C4 | 154.7 | |  | |  | |  |
|  | C5 | 121.3 | | 7.76, s | | Thz4-(C2, C4), Pyr-(C6) | |  |
| Ind | C=O | 161.4 | |  | |  | |  |
|  | C2 | 127.1 | |  | |  | |  |
|  | C3 | 109.8 | |  | |  | |  |
|  | C3a | 119.6 | |  | |  | |  |
|  | C3b | 65.9 | | H3ba: 4.22, d, *J* = 10.5  H3bb: 4.95, d, *J* = 10.5 | | Ind-(C2, C3, C3a), Glu-(C3)  Ind-(C2, C3, C3a) | | Glu-(H3), Ind-(H3bb, H4ab)  Glu-(H3), Ind-(H3ba), Ser-(NH) |
|  | C4 | 127.9 | |  | |  | |  |
|  | C4a | 68.4 | | H4aa: 5.00, d, *J* = 12.5  H4ab: 5.97, d, *J* = 12.5 | | Ind-(C3a, C4, C5), Glu-(C=O)  Ind-(C3a, C4, C5), Glu-(C=O) | | Ind-(H3ba, H4ab, H5)  Ind-(H3ba, H4aa, H5) |
|  | C5 | 123.6 | | 7.14, d, *J* = 7.0 | | Ind-(C3a, C4a, C6, C7) | | Ind-(H4aa/b, H6) |
|  | C6 | 125.1 | | 7.39, dd, *J* = 7.0, 8.5 | | Ind-(C4, C5, C7, C7a) | | Ind-(H5, H7) |
|  | C7 | 112.2 | | 7.80, d, *J* = 8.5 | | Ind-(C3a, C5) | | Ind-(H6) |
|  | C7a | 135.3 | |  | |  | |  |
|  | N-OH |  | | 10.51, s | | Ind-(C2) | |  |
| Sug1 | C1 | 101.8 | | 5.45, brs | | Thz3-(C5), Sug1-(C2, C3, C5) | | Sug1-(H2, H4, C2-OCH_3_) |
|  | C2 | 75.9 | | 4.04, m | | Sug1-(C1, C3, C4, C2-OMe) | | Sug1-(H1, H3, C2-OCH_3_) |
|  | C2-OMe | 59.4 | | 3.51, s | | Sug1-(C2) | | Sug1-(H1, H2) |
|  | C3 | 80.1 | | 3.64, m | | Sug1-(C4, C3-OMe) | | Sug1-(H2, C3-OCH_3_) |
|  | C3-OMe | 57.9 | | 3.45, s | | Sug1-(C3) | | Sug1-(H2, H4) |
|  | C4 | 77.3 | | 3.67, m | | Sug1-(C5), Sugar 2-(C1) | | Sug1-(H6, C3-OCH_3_), Sug2-(H1) |
|  | C5 | 70.0 | | 3.68, m | | Sug1-(C1, C3, C6) | |  |
|  | C6 | 17.4 | | 1.34, d, *J* = 5.5 | | Sug1-(C4, C5) | | Sug1-(H4) |
| Sug2 | C1 | 100.2 | | 4.71, dd, *J* = 1.5, 9.0 | | Sug1-(C4), Sug2-(C2) | | Sug1-(H4), Sug2-(H2b, H3, H5) |
|  | C2 | 38.3 | | H2a: 1.41, m  H2b: 2.20, m | | Sug2-(C1, C3, C4) | | Sug2-(H2b, H4)  Sug2-(H1, H2a, H3) |
|  | C3 | 69.6 | | 3.52, m | | Sug2-(C4) | | Sug2-(H1, H2b) |
|  | C4 | 88.2 | | 2.90, m | | Sug2-(C5, C6), Sug3-(C1) | | Sug3-(H1), Sug2-(H6) |
|  | C5 | 70.0 | | 3.23, m | | Sug2-(C1, C4, C6) | | Sug2-(H1, H3, H6) |
|  | C6 | 17.6 | | 1.18, d, *J* = 6.0 | | Sug2-(C4, C5) | | Sug2-(H4, H5) |
| Sug3 | C1 | 101.0 | | 4.45, dd, *J* = 2.0, 10.0 | | Sug2-(C4), Sug3-(C2, C3, C5) | | Sug2-(H4, H6), Sug3-(H2b, H3, H5) |
|  | C2 | 38.7 | | H2a: 1.56, m  H2b: 2.16, m | | Sug3-(C1, C3, C4) | | Sug3-(H2b, H4)  Sug3-(H1, H2b) |
|  | C3 | 70.9 | | 3.48, *ol* | |  | | Sug3-(H1, H2b, H5) |
|  | C4 | 76.7 | | 2.96, m | | Sug3-(C3, C5, C6) | | Sug3-(H2a, H6) |
|  | C5 | 72.3 | | 3.28, m | | Sug3-(C1, C3, C4, C6) | | Sug3-(H1, H6) |
|  | C6 | 17.4 | | 1.26, d, *J* = 6.0 | | Sug3-(C4, C5) | | Sug3-(H4, H5) |
| Sug4 | C1 | 100.5 | | 5.18, d, *J* = 7.5 | | Pyr-(C3), Sug4-(C3, C5) | | Pyr-(H4), Sug4-(H3, H5) |
|  | C2 | 69.5 | | 4.19, dd, *J* = 7.5, 9.5 | | Sug4-(C1, C3) | | Sug4-(H1, H3) |
|  | C3 | 84.3 | | 3.24, *ol* | | Sug4-(C1, C2, C3-OMe) | | Sug4-(H1, H2, H5) |
|  | C3-OMe | 58.1 | | 3.50, s | | Sug4-(C3) | | Sug4-(H3) |
|  | C4 | 77.6 | | 3.48, *ol* | | Sug4-(C4-OMe) | | Sug4-(H5, H6) |
|  | C4-OMe | 61.9 | | 3.56, s | | Sug4-(C4) | | Sug4-(H6) |
|  | C5 | 71.3 | | 3.78, m | | Sug4-(C1, C3, C4, C6) | | Sug4-(H1, H3, H4, H6) |
|  | C6 | 16.8 | | 1.35, d, *J* = 6.5 | | Sug4-(C4, C5) | | Sug4-(H4, H5, C4-OCH_3_) |

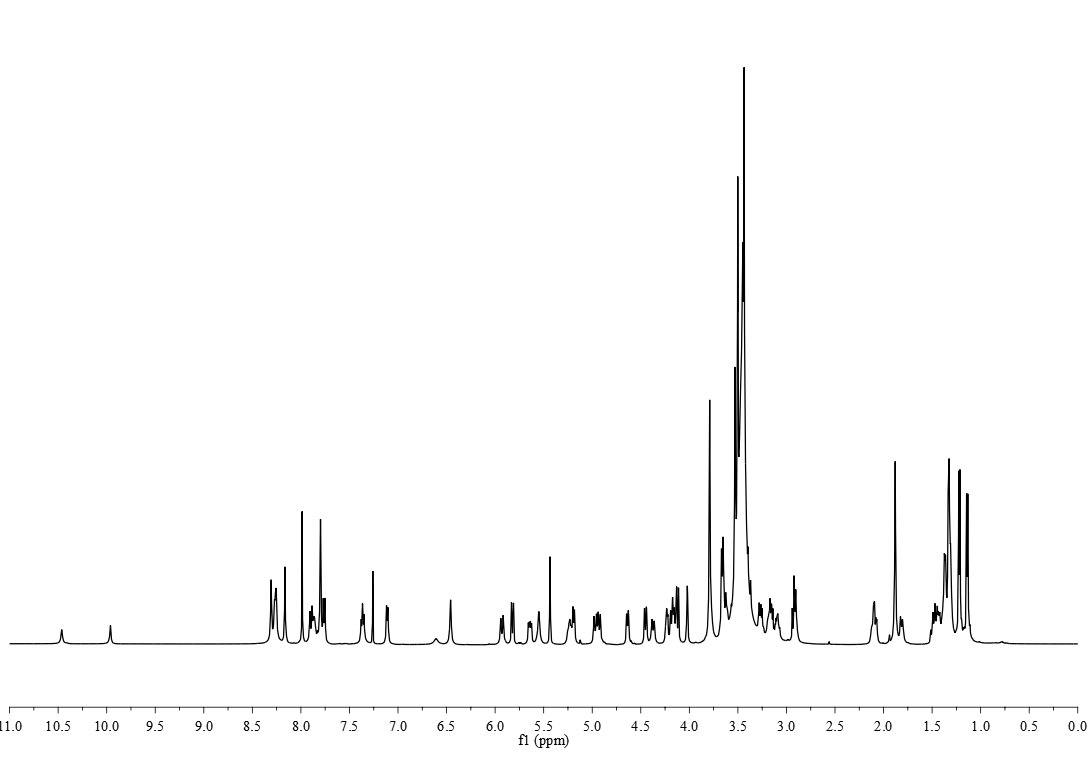

**Figure S3.** ^1^H NMR spectrum (500 MHz) of persiathiacin A in CDCl_3_-CD_3_OD (9:1).

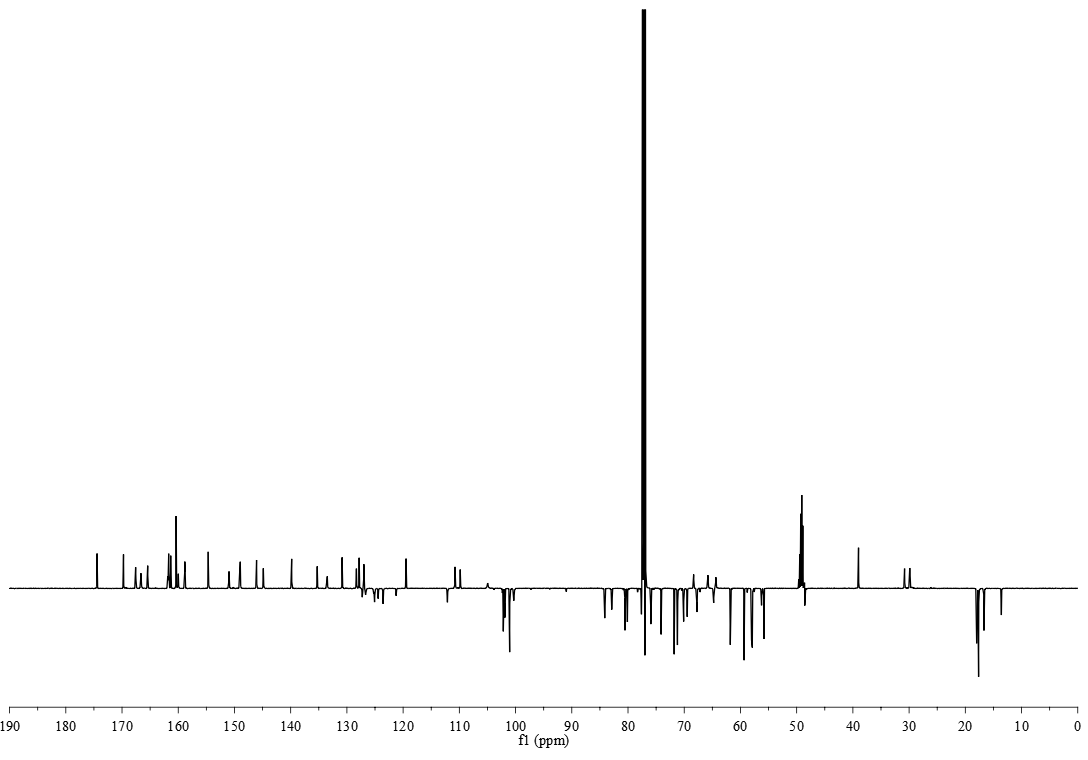

**Figure S4.** ^13^C NMR spectrum (125 MHz) of persiathiacin A in CDCl_3_-CD_3_OD (9:1).

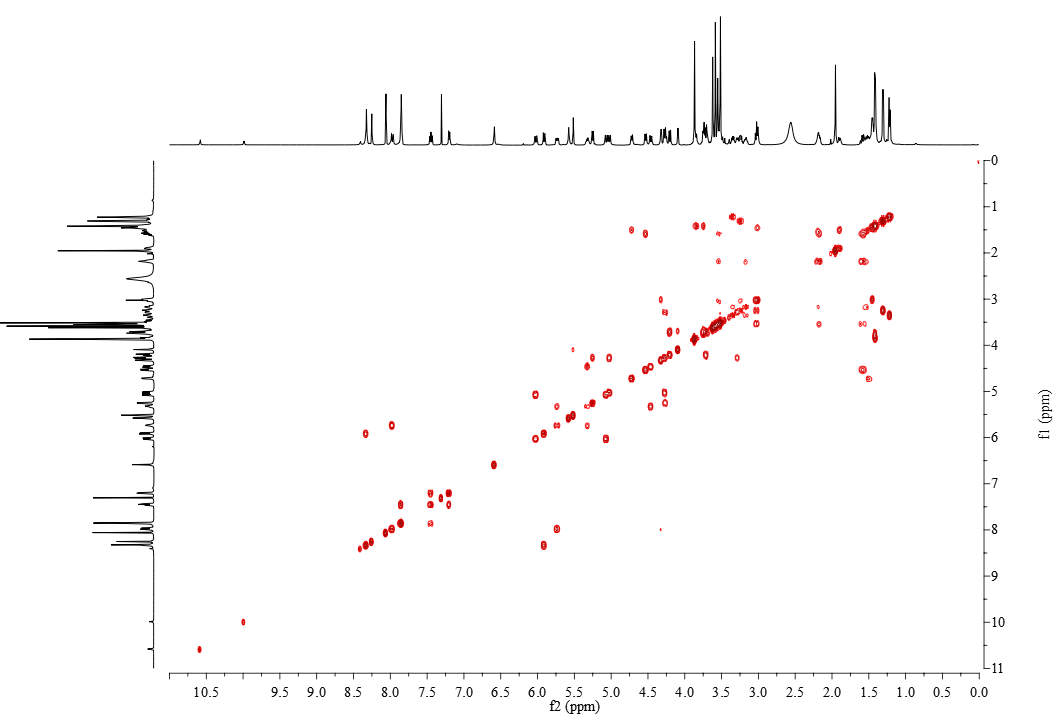

**Figure S5.** ^1^H-^1^H COSY spectrum of persiathiacin A in CDCl_3_-CD_3_OD (9:1).

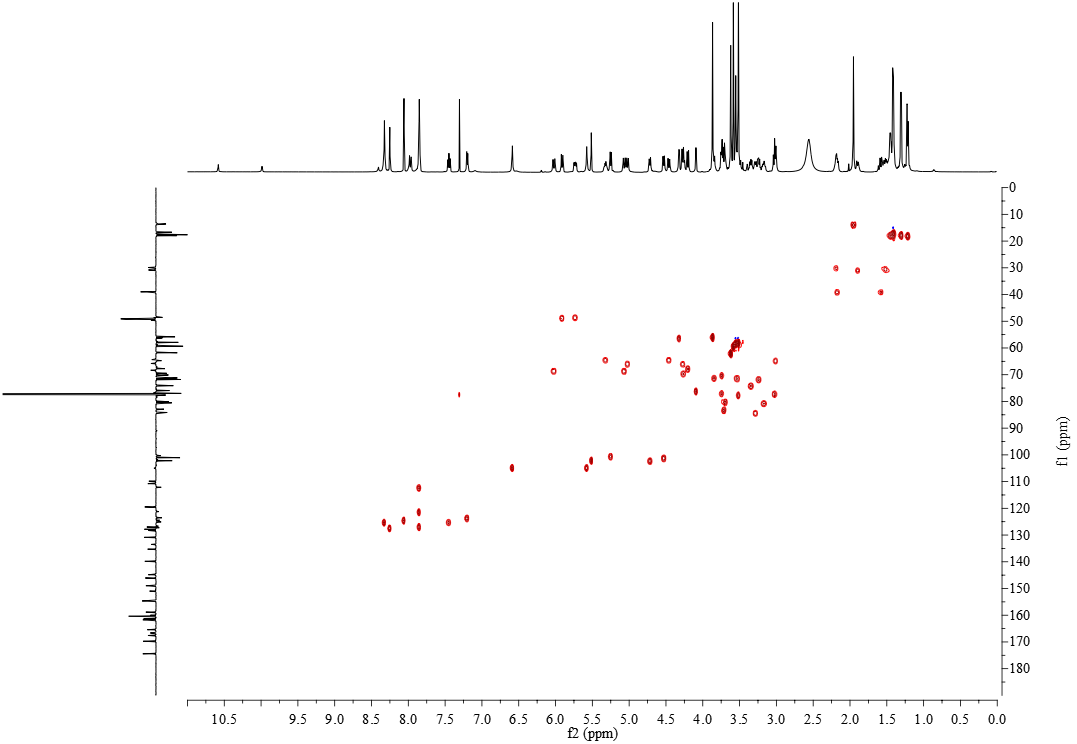

**Figure S6.** ^1^H-^13^C HSQC spectrum of persiathiacin A in CDCl_3_-CD_3_OD (9:1).

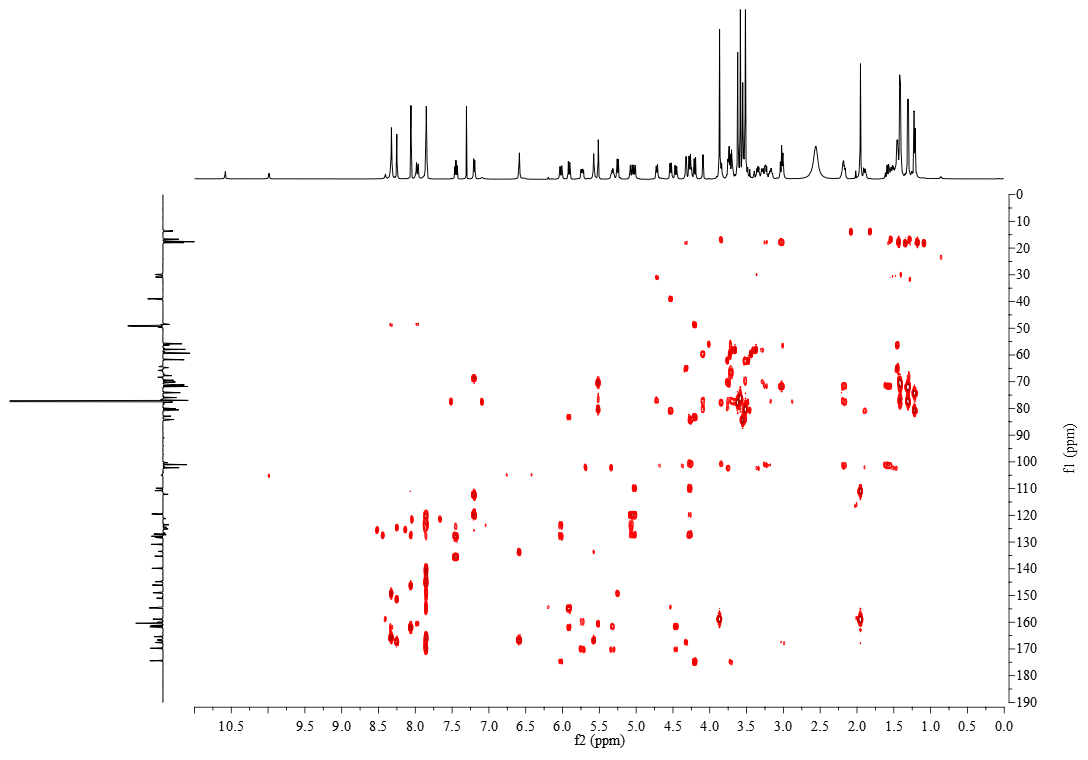

**Figure S7.** ^1^H-^13^C HMBC spectrum of persiathiacin A in CDCl_3_-CD_3_OD (9:1).

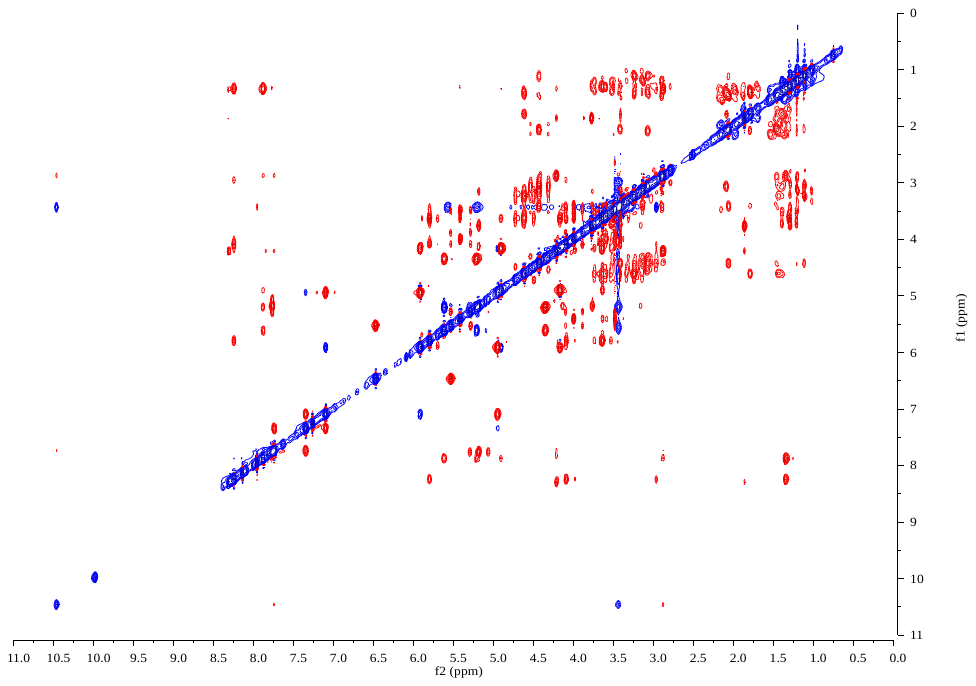

**Figure S8.** ^1^H-^1^H ROESY spectrum of persiathiacin A in CDCl_3_-CD_3_OD (9:1).

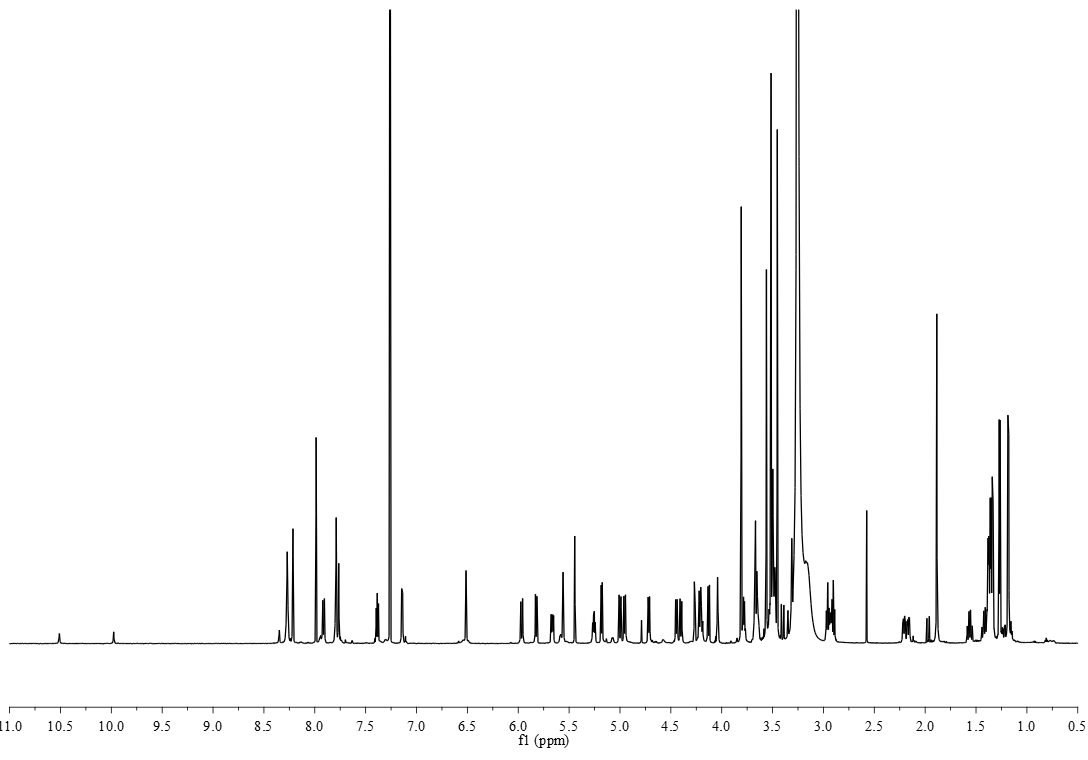

**Figure S9.** ^1^H NMR spectrum (700 MHz) of persiathiacin B in CDCl_3_-CD_3_OD (9:1).

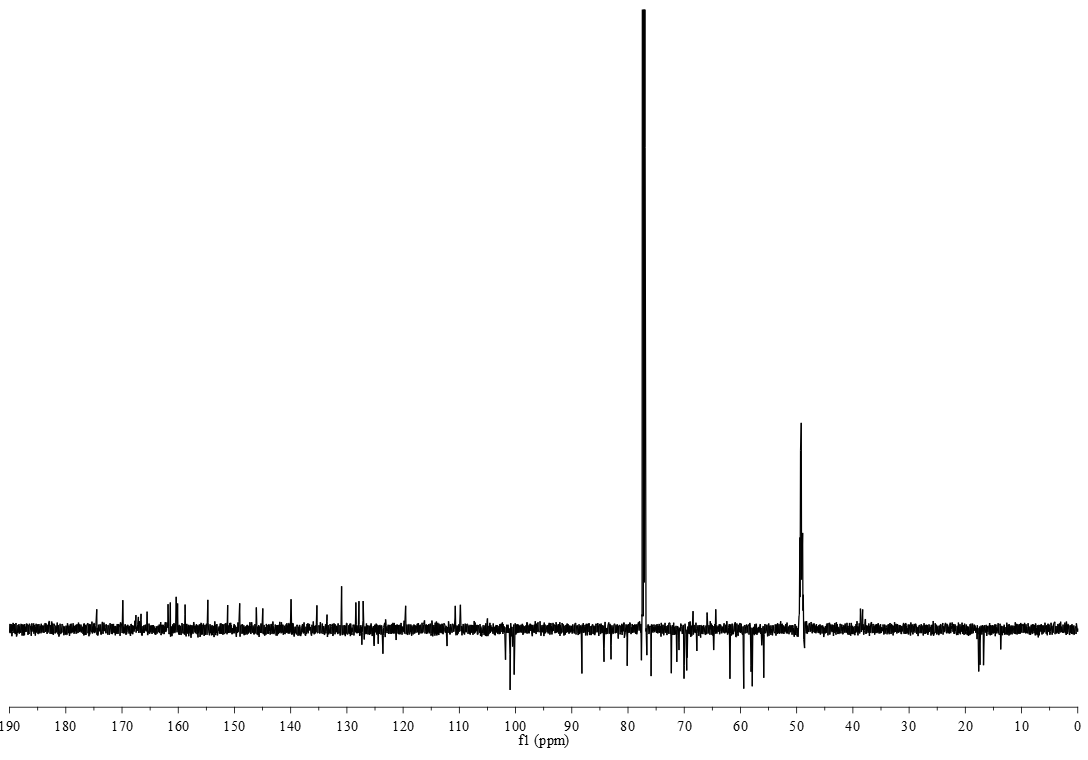

**Figure S10.** ^13^C NMR spectrum (176 MHz) of persiathiacin B in CDCl_3_-CD_3_OD (9:1).

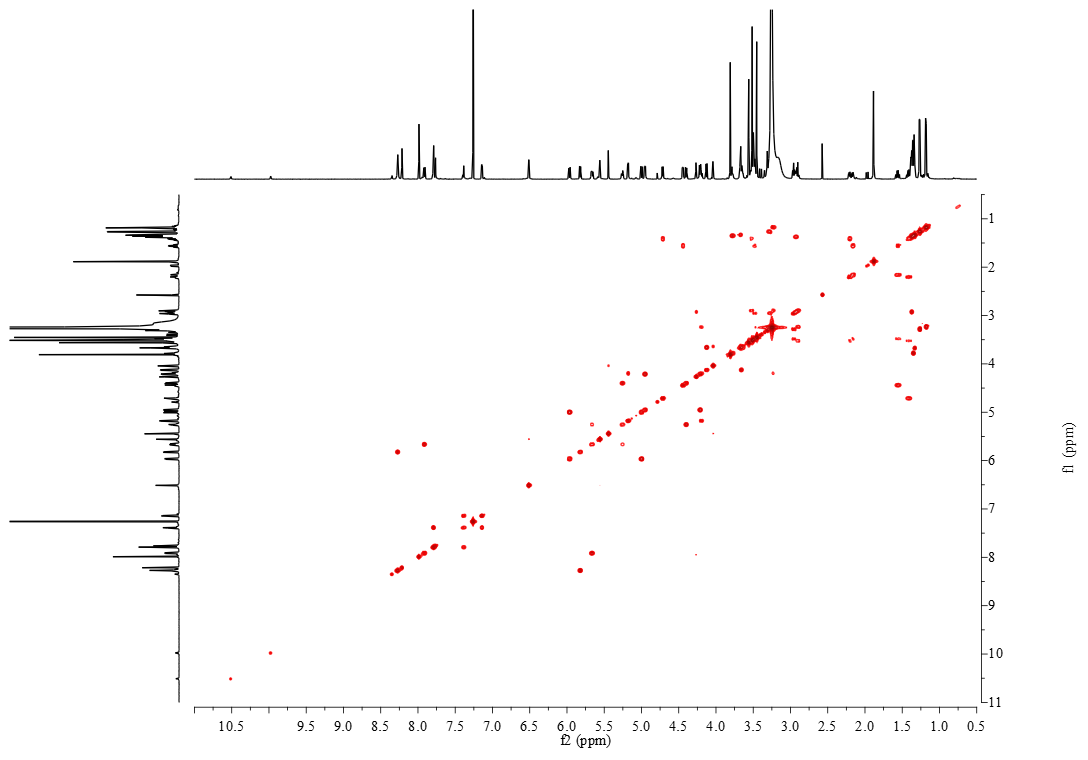

**Figure S11.** ^1^H-^1^H COSY spectrum of persiathiacin B in CDCl_3_-CD_3_OD (9:1).

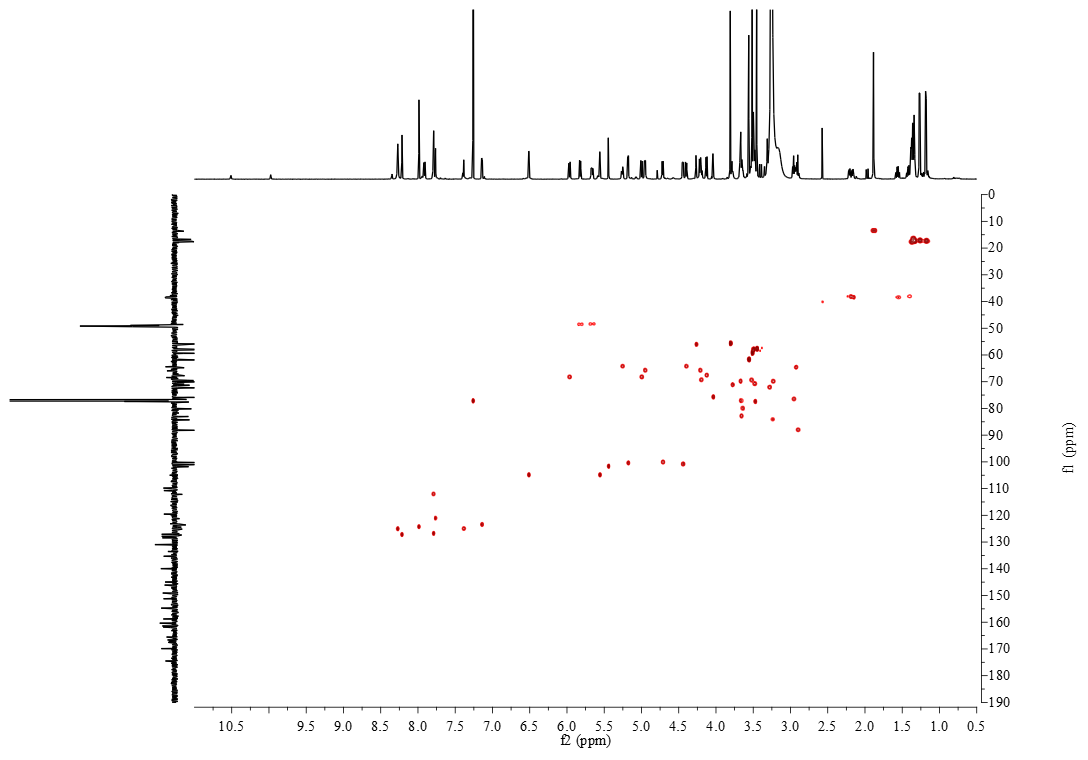

**Figure S12.** ^1^H-^13^C HSQC spectrum of persiathiacin B in CDCl_3_-CD_3_OD (9:1).

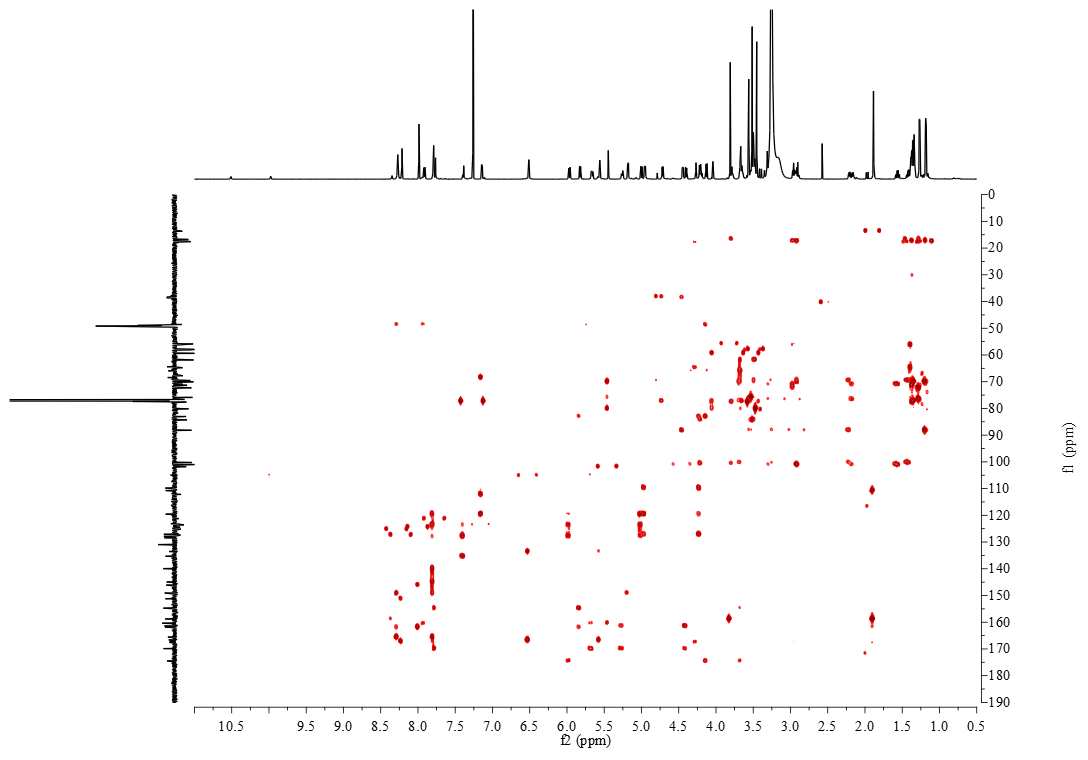

**Figure S13.** ^1^H-^13^C HMBC spectrum of persiathiacin B in CDCl_3_-CD_3_OD (9:1).

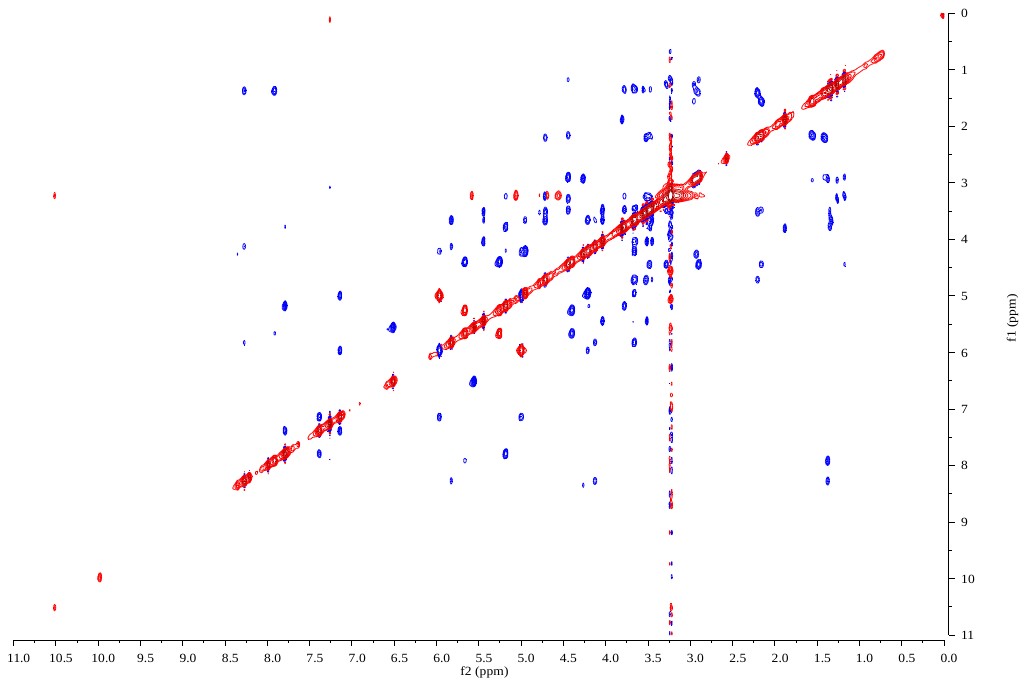

**Figure S14.** ^1^H-^1^H ROESY spectrum of persiathiacin B in CDCl_3_-CD_3_OD (9:1).

**Table S3.** Predicted secondary metabolite biosynthetic gene clusters in the genome of *Actinokineospora* sp. UTMC 2448.

| **Cluster** | **antiSMASH functional prediction** | **Start position** | **End position** |
| --- | --- | --- | --- |
| Cluster 1 | [Arylpolyene](http://antismash.secondarymetabolites.org/help#arylpolyene) | 361740 | 403998 |
| Cluster 2 | [Ectoine](http://antismash.secondarymetabolites.org/help#bacteriocin) | 427620 | 438018 |
| Cluster 3 | Terpene | 557729 | 606490 |
| Cluster 4 | [T1pks](http://antismash.secondarymetabolites.org/help#t1pks)-[Nrps](http://antismash.secondarymetabolites.org/help#nrps) | 830346 | 897979 |
| Cluster 5 | Terpene | 1663302 | 1684207 |
| Cluster 6 | Bacteriocin | 1814304 | 1825113 |
| Cluster 7 | [Siderophore](http://antismash.secondarymetabolites.org/help#siderophore) | 1872946 | 1884691 |
| Cluster 8 | [Nrps](http://antismash.secondarymetabolites.org/help#nrps)-[Lantipeptide](http://antismash.secondarymetabolites.org/help#lantipeptide) | 1895748 | 1950324 |
| Cluster 9 | [T1pks](http://antismash.secondarymetabolites.org/help#t1pks) | 1951254 | 1997820 |
| Cluster 10 | [Terpene](http://antismash.secondarymetabolites.org/help#terpene) | 2341218 | 2362180 |
| Cluster 11 | [Thiopeptide](http://antismash.secondarymetabolites.org/help#thiopeptide)-[Oligosaccharide](http://antismash.secondarymetabolites.org/help#oligosaccharide) | 2544416 | 2605135 |
| Cluster 12 | [T2pks](http://antismash.secondarymetabolites.org/help#t2pks) | 2622827 | 2665306 |
| Cluster 13 | [T3pks](http://antismash.secondarymetabolites.org/help#t3pks) | 2694719 | 2735756 |
| Cluster 14 | [Otherks](http://antismash.secondarymetabolites.org/help#otherks) | 2799423 | 2846466 |
| Cluster 15 | [T1pks](http://antismash.secondarymetabolites.org/help#t1pks) | 2851965 | 2898111 |
| Cluster 16 | [Indole](http://antismash.secondarymetabolites.org/help#indole) | 2927064 | 2948149 |
| Cluster 17 | [Phenazine](http://antismash.secondarymetabolites.org/help#phenazine) | 3024546 | 3045010 |
| Cluster 18 | [Nrps](http://antismash.secondarymetabolites.org/help#nrps) | 3156780 | 3256358 |
| Cluster 19 | [Terpene](http://antismash.secondarymetabolites.org/help#terpene) | 3260280 | 3281989 |
| Cluster 20 | [Nrps](http://antismash.secondarymetabolites.org/help#nrps) | 3330279 | 3386499 |
| Cluster 21 | [Ectoine](http://antismash.secondarymetabolites.org/help#ectoine) | 3571813 | 3582196 |
| Cluster 22 | [Butyrolactone](http://antismash.secondarymetabolites.org/help#butyrolactone)-[Nrps](http://antismash.secondarymetabolites.org/help#nrps) | 3579882 | 3652648 |
| Cluster 23 | [T1pks](http://antismash.secondarymetabolites.org/help#t1pks) | 3904090 | 3957797 |
| Cluster 24 | [Other](http://antismash.secondarymetabolites.org/help#other) | 4140452 | 4181138 |
| Cluster 25 | [Terpene](http://antismash.secondarymetabolites.org/help#terpene) | 4239895 | 4261055 |
| Cluster 26 | [Melanin](http://antismash.secondarymetabolites.org/help#melanin) | 4434945 | 4445863 |
| Cluster 27 | [T1pks](http://antismash.secondarymetabolites.org/help#t1pks) | 4577600 | 4623818 |
| Cluster 28 | [Lantipeptide](http://antismash.secondarymetabolites.org/help#lantipeptide) | 4650662 | 4674898 |
| Cluster 29 | [Ladderane](http://antismash.secondarymetabolites.org/help#ladderane)-[Arylpolyene](http://antismash.secondarymetabolites.org/help#arylpolyene)-[Nrps](http://antismash.secondarymetabolites.org/help#nrps) | 4820757 | 4897663 |
| Cluster 30 | [Lantipeptide](http://antismash.secondarymetabolites.org/help#lantipeptide)-[T1pks](http://antismash.secondarymetabolites.org/help#t1pks)-NRPS | 5700170 | 5876737 |
| Cluster 31 | Bacteriocin | 6203188 | 6214081 |
| Cluster 32 | [T1pks](http://antismash.secondarymetabolites.org/help#t1pks)-[Butyrolactone](http://antismash.secondarymetabolites.org/help#butyrolactone) | 6252778 | 6312920 |

**Table S4.** Genes in the persiathiacin biosynthetic gene cluster and the percentage identity of the proteins they encode to proteins of known function.

| **Gene/Protein** | **Locus_tag** | **Length**  **bp/aa** | **Similar Proteins** | **% aa Identity** |
| --- | --- | --- | --- | --- |
| *perO*/PerO | Actin_02399 | 906/302 | NocO (*Nocardia* sp. ATCC202099)  NosO (*Streptomyces actuosus*) | 52  44 |
| *perN*/PerN | Actin_02400 | 1236/412 | NocN (*Nocardia* sp. ATCC202099)  NosN (*Streptomyces actuosus*) | 85  75 |
| *perM*/PerM | Actin_02401 | 147/49 | NocM (*Nocardia* sp. ATCC202099)  NosM (*Streptomyces actuosus*) | 98  88 |
| *perL*/PerL | Actin_02402 | 1188/396 | NocL (*Nocardia* sp. ATCC202099)  NosL (*Streptomyces actuosus*) | 85  82 |
| *perK*/PerK | Actin_02403 | 846/282 | NocK (*Nocardia* sp. ATCC202099)  NosK (*Streptomyces actuosus*) | 67  59 |
| *perI*/PerI | Actin_02404 | 1260/420 | NocI (*Nocardia* sp. ATCC202099)  NosI (*Streptomyces actuosus*) | 67  62 |
| *perH*/PerH | Actin_02405 | 1728/576 | NocH (*Nocardia* sp. ATCC202099)  NosH (*Streptomyces actuosus*) | 54  49 |
| *perG*/PerG | Actin_02406 | 1854/618 | NocG (*Nocardia* sp. ATCC202099)  NosG (*Streptomyces actuosus*) | 73  65 |
| *perF*/PerF | Actin_02407 | 1398/466 | NocF (*Nocardia* sp. ATCC202099)  NosF (*Streptomyces actuosus*) | 47  44 |
| *perE*/PerE | Actin_02408 | 2589/863 | NocE (*Nocardia* sp. ATCC202099)  NosE (*Streptomyces actuosus*) | 64  56 |
| *perD*/PerD | Actin_02409 | 966/322 | NocD (*Nocardia* sp. ATCC202099)  NosD (*Streptomyces actuosus*) | 59  55 |
| *perC*/PerC | Actin_02410 | 1221/407 | NocC (*Nocardia* sp. ATCC202099)  NosC (*Streptomyces actuosus*) | 81  71 |
| *perV*/PerV | Actin_02411 | 1122/374 | NocV (*Nocardia* sp. ATCC202099) | 71 |
| *perR*/PerR | Actin_02412 | 516/172 | NocR (*Nocardia* sp. ATCC202099) | 87 |
| *perQ*/PerQ | Actin_02413 | 597/199 | NocQ (*Nocardia* sp. ATCC202099) | 78 |
| *perU*/PerU | Actin_02414 | 1173/391 | NocU (*Nocardia* sp. ATCC202099) | 59 |
| *perT*/PerT | Actin_02415 | 1122/374 | NocT (*Nocardia* sp. ATCC202099) | 59 |
| *perS1*/PerS1 | Actin_02416 | 1020/340 | Oxidoreductase (*Streptomyces* sp.)  NDP-hexose-3-ketoreductase (*Streptoalloteichus hindustanus*) | 61  59 |
| *perP*/PerP | Actin_02417 | 993/331 | NocP (*Nocardia* sp. ATCC202099)  NosP (*Streptomyces actuosus*) | 61  55 |
| *perB*/PerB | Actin_02418 | 1185/395 | NosB (*Streptomyces actuosus*)  NocB (*Nocardia* sp. ATCC202099) | 55  54 |
| *perS2*/PerS2 | Actin_02419 | 858/286 | UDP-glucose 4-epimerase (*Amycolatopsis pretoriensis*)  SpeI (*Streptomyces spectabilis*) | 44  43 |
| *perS3*/PerS3 | Actin_02420 | 819/273 | macrocin O-methyltransferase (*Micromonospora viridifaciens*)  putative methyl transferase (*Streptomyces argenteolus*) | 60  51 |
| *perS4*/PerS4 | Actin_02421 | 1137/379 | rhamnosyltransferase (*Streptomyces* sp. SANK 60405)  glycosyltransferase (*Allokutzneria albata*) | 40  39 |
| *perS5*/PerS5 | Actin_02422 | 786/262 | class I SAM-dependent methyltransferase  (*Kitasatospora mediocidica*)  (Micromonospora viridifaciens) | 53  51 |
| *perX*/PerX | Actin_02423 | 1176/392 | cytochrome P450  (*Actinobacteria bacterium* OK074)  (*Streptomyces gilvigriseus*) | 41  40 |
| *perS6*/PerS6 | Actin_02424 | 1197/399 | glycosyltransferase  (*Streptomyces aizunensis*)  (*Kutzneria* sp. 744) | 45  41 |
| *perS7*/PerS7 | Actin_02425 | 1134/378 | class I SAM-dependent methyltransferase (*Actinoalloteichus hymeniacidonis*)  (*Nocardia puris*) | 70  52 |
| *perS8*/PerS8 | Actin_02426 | 1044/348 | glycosyltransferase  (*Streptomyces aizunensis*)  (*Streptomyces* sp. WMMB 714) | 50  45 |
| *perS9*/PerS9 | Actin_02427 | 1086/362 | glycosyltransferase  (*Amycolatopsis tolypomycina*)  (*Streptomyces* sp. PAN_FS17) | 41  40 |
| *perS10*/PerS10 | Actin_02428 | 1404/468 | NDP-hexose 2,3-dehydratase (*Plantactinospora* sp. KBS50)  (*Amycolatopsis vancoresmycina*) | 60  59 |
| *perS11*/PerS11 | Actin_02429 | 912/304 | NDP-hexose 4-ketoreductase (*Micromonospora* sp. CNZ309)  (*Micromonospora echinofusca*) | 56  56 |
| *perS12*/PerS12 | Actin_02430 | 1305/435 | NDP-hexose 3,4-dehydratase (*Streptomyces minoensis*)  (*Streptomyces caniferus*) | 71  70 |
| *perA*/PerA | Actin_02431 | 435/145 | NocA (*Nocardia* sp. ATCC202099)  NosA (*Streptomyces actuosus*) | 59  67 |

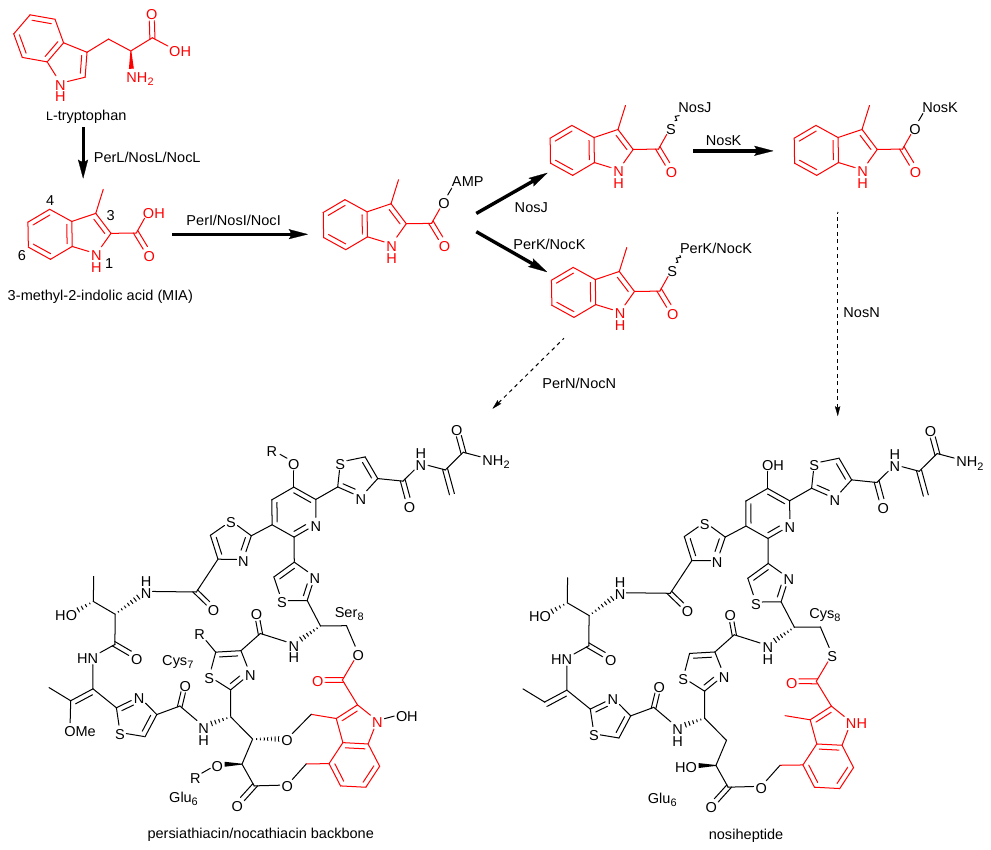

**Figure S15.** Similarities and differences in the activation and attachment of MIA to the nosiheptide, persiathiacin, and nocathiacin processed core peptides (R-groups in persiathiacin and nocathiacin are shown in figures 1 and 2 of the main manuscript).

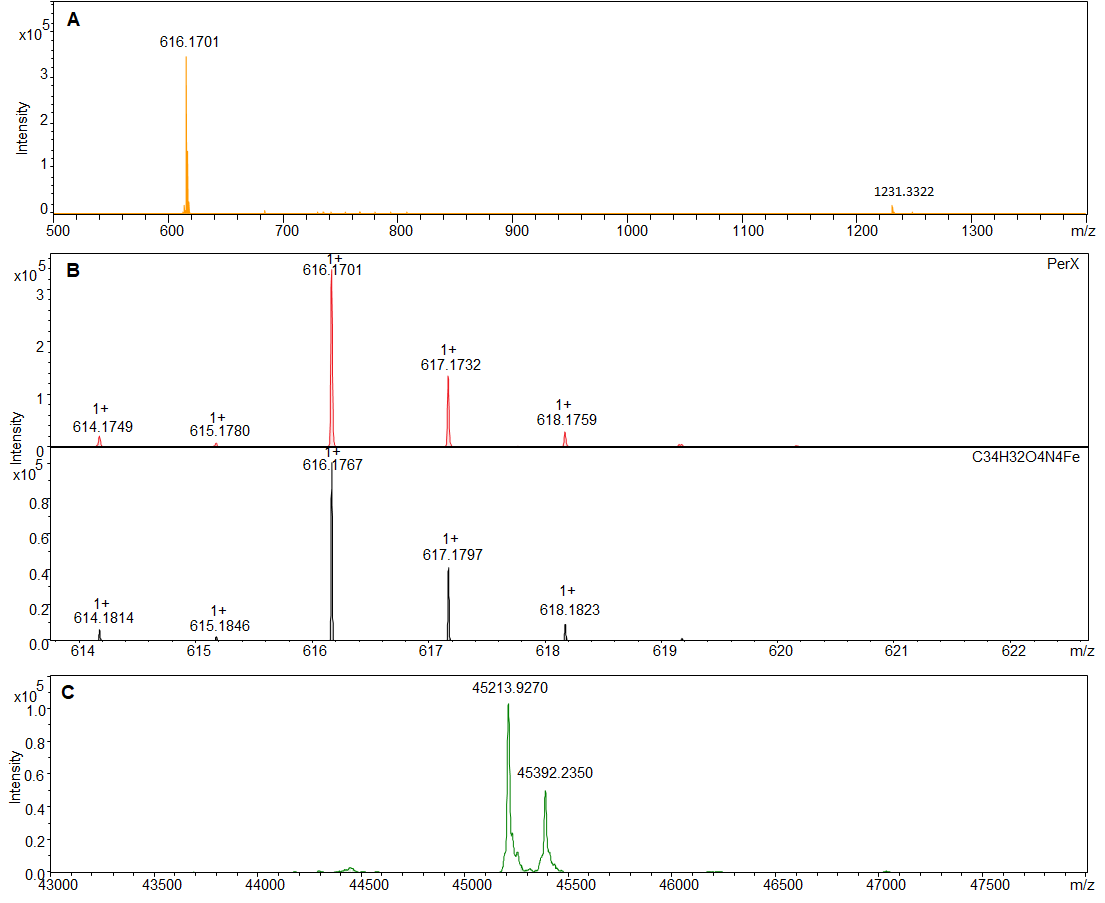

**Figure S16.** ESI-Q-TOF-MS analysis of purified recombinant PerX. (A) An ion with *m/z* = 6161.1701, corresponding to [M]^+^ for ferric haem is observed. (B) Comparison of the measured (top) and simulated (bottom) spectra for the [M]^+^ ion of ferric haem. (C) Deconvoluted mass spectrum of PerX. The measured masses correspond to the protein after loss of the N-terminal methionine residue (measured: 45213.92, calculated: 45215.35) and a glucuronidated derivative (measured: 45392.23, calculated: 45391.35).

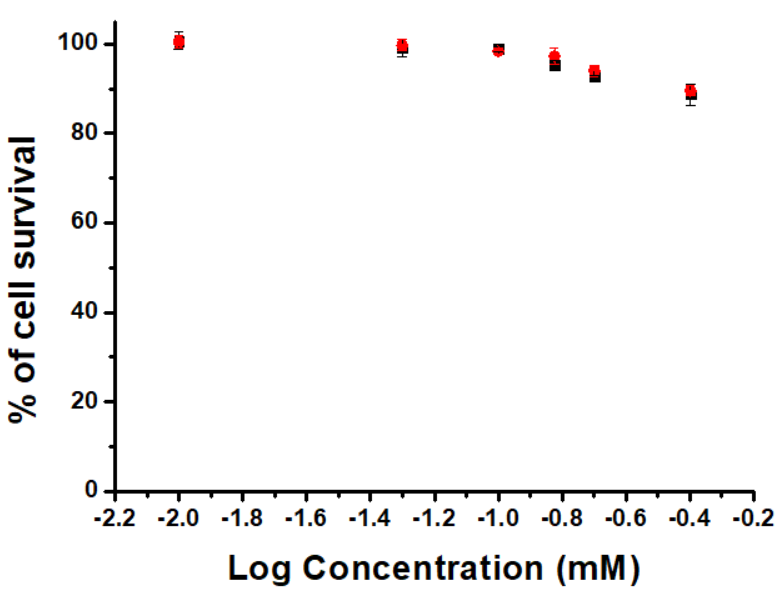

**Figure S17.** Percentage of A2780 ovarian cancer cells surviving after exposure to various concentration of persiathiacin A ranging from 10 to 400 μM relative to untreated (vehicle) controls. These experiments included 24 h of drug exposure time and 72 h of recovery time in drug free medium.

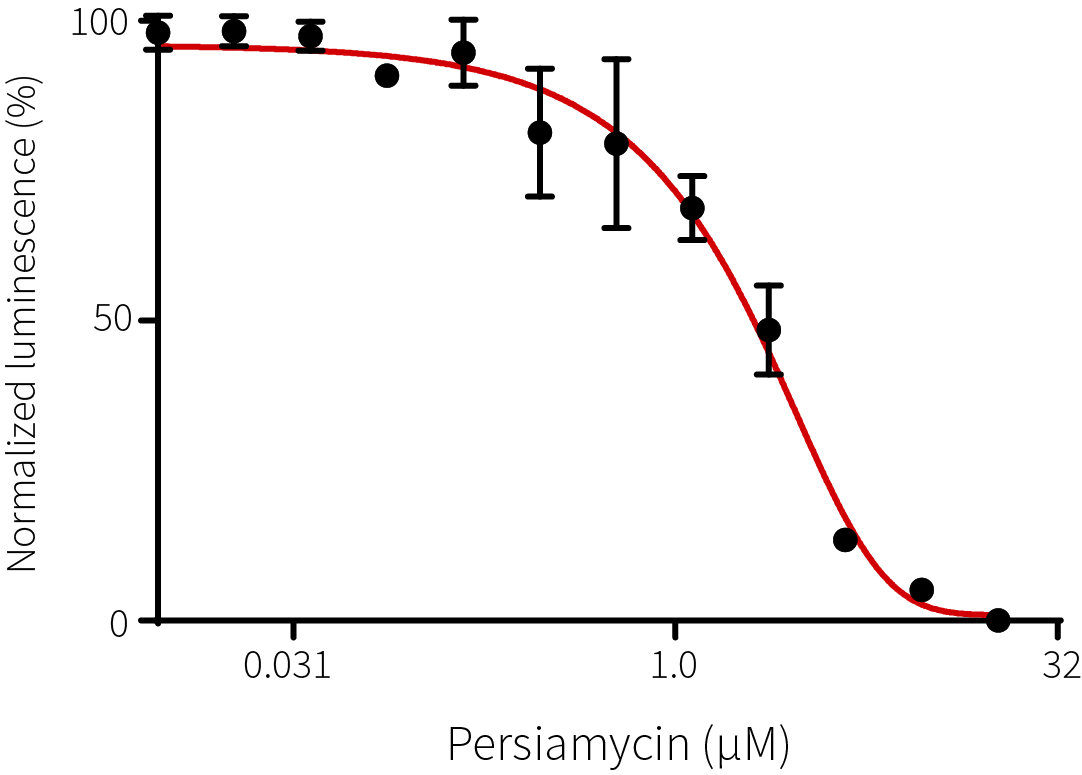

**Figure S18.** Concentration-dependent inhibition of the *E. coli* ribosome by persiathiacin A.
